# Gene expression in the cardiovascular system of the domestic sheep (*Ovis aries*); a new tool to advance our understanding of cardiovascular disease

**DOI:** 10.1101/2020.04.24.059857

**Authors:** Hiu-Gwen Tsang, Emily L. Clark, Greg R. Markby, Stephen J. Bush, David A. Hume, Brendan M. Corcoran, Vicky E. MacRae, Kim M. Summers

**Author notes:** These authors contributed equally to this work. These authors contributed equally to this work (Hiu-Gwen Tsang, Emily L. Clark).

## Abstract

Large animal models are of increasing importance in cardiovascular disease research as they demonstrate more similar cardiovascular features (in terms of anatomy, physiology and size) to humans than do rodent species. The maintenance of a healthy cardiovascular system requires expression of genes that contribute to essential biological activities and repression of those that are associated with functions likely to be detrimental to cardiovascular homeostasis. In this study we have used the transcriptome of the sheep, which has been utilised extensively to model human physiology and disease, to explore genes implicated in the process of vascular calcification. Vascular calcification is a major disruption to cardiovascular homeostasis where tissues of the cardiovascular system undergo ectopic calcification and consequent dysfunction. We investigate the gene expression profiles of genes involved in vascular calcification in a wide array of cardiovascular tissues and across multiple developmental stages, using RT-qPCR. The majority of transcriptomic studies on the mammalian cardiovascular system to date have focused on regional expression of specific genes. Here we also use RNA sequencing results from the sheep heart and cardiac valves to further explore the transcriptome of the cardiovascular system in this large animal. Our results demonstrate that there is a balance between genes that promote and those that suppress mineralisation during development and across cardiovascular tissues. We show extensive expression of genes encoding proteins involved in formation and maintenance of the extracellular matrix in cardiovascular tissues, and high expression of haematopoietic genes in the cardiac valves. Our analysis will support future research into the functions of implicated genes in the development of vascular calcification, and increase the utility of the sheep as a large animal model for understanding cardiovascular disease. This study provides a foundation to explore the transcriptome of the developing cardiovascular system and is a valuable resource for the fields of mammalian genomics and cardiovascular research.

## Introduction

The cardiovascular system plays a crucial role not only in the distribution of nutrients to the various cells, tissues and organs within the mammalian body, but also in the removal of waste products. Extensive regulatory mechanisms are required to support this functional system, with perturbations likely to lead to abnormalities, and thus give rise to cardiovascular-related diseases. Cardiovascular disease (CVD) is a major cause of morbidity and mortality worldwide, with an estimated 17.3 to 17.5 million deaths per year (Townsend et al. 2016; WHO, 2017). The major cardiovascular-related causes of premature death include coronary heart disease (CHD) and stroke (Townsend et al. 2016; WHO, 2017). In addition, cardiac valvulopathies are becoming increasingly prevalent in the ageing population (Nkomo et al. 2006). A recent United Kingdom study of nearly 80,000 adult patients referred for echocardiography found that 50% had some degree of cardiac valve dysfunction (Marciniak et al. 2017). In contract, only 12% of the same patient group had left ventricular systolic dysfunction. Aortic aneurysm is also an increasing health care burden, particularly in elderly males (reviewed by Hohneck et al, 2019) who found an incidence of 7% in elderly male patients hospitalised for cardiopulmonary symptoms). Thus cardiovascular disease is a major socioeconomic burden and understanding of the cardiovascular system is important in developing treatments for the range of conditions.

In recent decades, both non-invasive and invasive CVD therapies have advanced considerably. This advancement has been underpinned by basic research, with animal models of CVD being of key importance. Of growing value is the use of large animal models of CVD research (reviewed in Tsang et al. 2016). Sheep and pigs, for example, are more similar in terms of their cardiovascular features (in terms of anatomy, genetics, physiology and size) to humans than are rodent species. Evidence suggests there are significant phenotypic differences between mouse and human stem cells (Ginis et al. 2004; Gabdoulline et al. 2015) and large animals might therefore provide greater similarity at the cellular and molecular level. Early developmental stages can be studied in detail in large animal models, which is limited in scope in both human and mouse (Emmert et al. 2013). Finer resolution of regions of the cardiovascular system is also possible with the increased size of the heart and vessels of the large animal models. Sufficient RNA for transcriptomic studies can be obtained from a single animal, so that inter-animal variability can be assessed. The major benefit of large animals in CVD clinical research remains however their application in the development of interventional technologies and implantable devices (reviewed in Tsang et al. 2016). Characterising the normal transcriptome of the healthy mammalian cardiovascular system will allow better understanding of the cellular changes induced by these treatments.

The maintenance of a healthy cardiovascular system requires expression of genes that contribute to essential biological activities and repression of those that are associated with functions likely to be detrimental to cardiovascular homeostasis. A major pathological process that disrupts cardiovascular homeostasis is vascular calcification (VC), which is associated with aging, hypertension and atherosclerosis (Abedin, Tintut, & Demer, 2004; Towler, 2008; Zhu, Mackenzie, Farquharson, & Macrae, 2012, Tsang et al. 2016). VC is a disease of abnormal mineral metabolism, involving the deposition of calcium phosphate, in the form of hydroxyapatite (HA), in cardiovascular tissues, most critically in the arteries and cardiac valves, and is a significant, independent risk factor of cardiovascular mortality (Giachelli, 2004; Li, Yang, & Giachelli, 2006; Zhu et al. 2012). Most individuals above 60 years of age have gradually enlarging calcium deposits in their major arteries (Allison, Criqui, & Wright, 2004; Demer & Tintut, 2008). VC is a highly regulated, active process involving a variety of signalling pathways, with evidence suggesting the involvement of mechanisms similarly observed in bone formation (Boström et al. 1993; Lanzer et al. 2014). However, the exact molecular basis underpinning the complex process of VC, particularly the dysregulated expression of genes involved in cardiovascular function, has yet to be fully defined.

A number of factors have been implicated in the phenotypic transition of vascular smooth muscle cells (VSMCs) into osteocytic-, osteoblastic- and chondrocytic-like cells (Yang et al. 2004; Li et al. 2006; Giachelli, 2009; Zhu et al. 2011; Zhu et al. 2012). These include osteochondrogenic markers, including *TNAP* (*ALPL* gene), osteopontin (also known as secreted phosphoprotein 1; *SPP1* gene), and the transcription factor *RUNX2* (Steitz et al. 2001; Rajamannan et al. 2003; Lomashvili et al. 2004; Speer et al. 2009; Yang et al. 2009; Zhu et al. 2012), as well as mineralisation inhibitors, including *ENPP1*, *MGP*, ecto-5’-nucleotidase *NT5E* (also known as cluster of differentiation 73, *CD73*), and *FBN1* (Luo et al. 1997; Schurgers et al. 2008; Rutsch et al. 2011; St Hilaire et al. 2011; Albright et al. 2015). During the calcification process VSMCs enter a synthetic state with abundant production of extracellular matrix (ECM) proteins (Hruska et al, 2005) followed by matrix vesicle-mediated calcification (Giachelli, 2009; Leopold, 2015). Indeed recent comparative transcription profiling has identified over 50 ECM genes identically regulated by calcifying VSMCs and bone-forming osteoblasts (Alves et al., 2014), with ECM proteins likely acting in concert with each other to determine the extent of calcification. Nevertheless, although the sequence of events conducting normal bone mineralisation is better understood, the specific mechanisms by which VC occurs remains ambiguous, as vascular cells may still retain their overall identity, despite acquiring osteoblastic properties (Frink, 2002; Zhu et al. 2012; Alves et al. 2014).

In recent years, the use of high-throughput technologies such as RNA sequencing (RNA-seq), has been continually expanding the number of gene expression datasets available for specific tissues and cells. In order to investigate the transcriptional landscape of the mammalian genome, various high-throughput mammalian gene expression profiling studies have been performed at different levels including varying cellular, tissue and whole organism levels. Atlases of gene expression have been generated for the majority of tissues, and a small number of cell types, in ovine (Jiang et al. 2014; Clark et al. 2017), caprine (Muriuki et al. 2019), bovine (Harhay et al. 2010), porcine (Freeman et al. 2012; Summers et al. 2020), equine (Mansour et al. 2017), murine, and human (Lein et al. 2007; Siddiqui et al. 2005; Su et al. 2002). Large-scale and collaborative projects have also been formed to generate big datasets on the mammalian transcriptome, across multiple tissues and cell types and including many individuals. These include the Functional Annotation of Animal Genomes (FAANG) Consortium (Andersson et al. 2015), Encyclopedia of DNA Elements (ENCODE) Consortium (Encode Project Consortium et al. 2007), the Functional ANnoTation Of the Mammalian genome (FANTOM5) consortium (Andersson et al. 2014; Fantom Consortium et al. 2014; Lizio et al. 2015), and the Genotype-Tissue Expression (GTEx) Consortium (Mele et al. 2015). The majority of transcriptomic studies on the mammalian cardiovascular system have to date only looked in detail at tissue-specific expression of single genes, or a subset of genes of interest (Gaborit et al. 2007; Potter, Abbey-Hosch, & Dickey, 2006), rather than providing wider resolution of the cardiovascular transcriptome across multiple tissues. Although there are various public resources, the transcriptomic data available for the mammalian cardiovascular system are generally limited to the “heart” or ventricular tissue, such as in BioGPS (http://biogps.org/) and the Expression Atlas online database (EMBL-EBI; https://www.ebi.ac.uk/gxa). In the human GTex Project (Mele et al. 2015), for example, two cardiovascular tissues are included (Heart – Atrial Appendage and Heart – Left Ventricle) from a large number of individuals (n=372 and n=386 respectively). RNA-seq has been used to greatly enhance resolution of cardiovascular disease in humans (reviewed in Wirka et al. 2018) and generate baseline estimates of gene expression in developing cardiovascular tissues (Pervolaraki et al. 2018). However, comparable resources were not available for large animal models, which could be used to develop interventions and other treatments for cardiovascular disease.

This study describes gene expression in the mammalian cardiovascular system, supporting and extending the high resolution gene expression atlas for sheep (Clark et al. 2017). We use reverse transcriptase quantitative PCR (RT-qPCR) to measure myocardial and arterial tissue gene expression during development in the sheep and investigate expression of vascular calcification (VC) inhibitors in the healthy cardiovascular system. We also present tissue-specific gene expression profiles in the heart muscle and cardiac valves using RNA-Seq. The study provides a foundation to explore the transcriptome of the developing cardiovascular system and provides a valuable resource for the fields of mammalian genomics and cardiovascular research.

## Materials and methods

### Reverse transcriptase quantitative PCR (RT-qPCR)

RT-qPCR was performed to measure the expression of twenty-five genes involved in cardiovascular and skeletal muscle function and VC across five developmental stages from Texel x Scottish Blackface sheep: 100-day gestation (foetal), newborn, 1 week, 8 weeks and 2 years (n=3-5 per group). Details of the samples included in each of the three sets of analyses (RNA-seq of eight tissues, developmental stage expression profiles and VC gene expression profiles) are included in Table 1. Samples were collected within an hour and thirty minutes post euthanasia. 17 different tissues were collected from adults; the equivalent tissues were collected from fetuses at day 100 of gestation and young lambs where possible. Detailed dissection of tissues was performed by the same two researchers, for all sheep, in order to standardise tissue sampling. After dissection, tissues of interest were placed into RNAlater (Thermo Fisher Scientific) and stored according to the manufacturer’s instructions. RNA was extracted from tissues using TRIzol (Thermo Fisher Scientific) as described in Clark et al. 2017. RNA integrity (RIN^e^) was estimated on an Agilent 2200 Tapestation System (Agilent Genomics) and only samples with RIN^e^ > 7 were included in the analysis. RT-qPCR reactions were performed using PrecisionPLUS-MX-SY Mastermix (containing SYBR Green; Primerdesign Ltd) following the manufacturer’s protocol. Details of the twenty-five sheep-specific primers used are listed in Supplementary Table 1. Primers were designed using the current version of the sheep genome Oar v3.1 (https://www.ensembl.org/Ovis_aries/Info/Annotation) with Primer3 software (http://primer3.ut.ee/) to span exon-exon junctions, and obtained from Invitrogen (Paisley, UK) and Primerdesign Ltd (Eastleigh, UK). Because of the limitations of the sheep genome sequence it was not possible to design primers for all genes of interest.

**Table 1.** Details of the samples, number of biological replicates and developmental stage of all samples included in the analyses.

| **Tissue** | **No. of replicates** | **Developmental stage** |
| --- | --- | --- |
| **RNA-seq** |  |  |
| **Aortic Valve** | 4 | Adult (2 years) |
| **Mitral Valve** | 4 | Adult (2 years) |
| **Tricuspid Valve** | 4 | Adult (2 years) |
| **Left Auricle** | 4 | Adult (2 years) |
| **Right Auricle** | 5 | Adult (2 years) |
| **Left Ventricle** | 6 | Adult (2 years) |
| **Right Ventricle** | 6 | Adult (2 years) |
| **Skeletal Muscle (Bicep)** | 6 | Adult (2 years) |
| **RT-qPCR (developmental stages)** |  |  |
| **Left Ventricle** | 3-5 | Fetus d100, newborn, 8 weeks, 2 years |
| **Interventricular septum** | 3-5 | Fetus d100, newborn, 1 week, 8 weeks, 2 years |
| **Aortic Root** | 3-5 | Newborn, 1 week, 8 weeks, 2 years |
| **Aortic Arch** | 3-5 | Newborn, one week, 2 years |
| **Abdominal Aorta** | 3-5 | Newborn, 8 weeks, 2 years |
| **Pulmonary Artery** | 3-5 | Newborn, 8 weeks, 2 years |
| **RT-qPCR (VC genes)** |  |  |
| **Left Auricle** | 6 | Adult (2 years) |
| **Left Atrium** | 6 | Adult (2 years) |
| **Right Auricle** | 6 | Adult (2 years) |
| **Right Atrium** | 6 | Adult (2 years) |
| **Right Ventricle** | 6 | Adult (2 years) |
| **Cranial Vena Cava** | 6 | Adult (2 years) |
| **Aortic Valve** | 6 | Adult (2 years) |
| **Mitral Valve** | 6 | Adult (2 years) |
| **Tricupsid Valve** | 6 | Adult (2 years) |
| **Pulmonary Valve** | 6 | Adult (2 years) |
| **Left Ventricle** | 6 | Adult (2 years) |
| **Interventricular septum** | 6 | Adult (2 years) |
| **Aortic Base** | 6 | Adult (2 years) |
| **Aortic Arch** | 6 | Adult (2 years) |
| **Descending Thoracic Aorta** | 6 | Adult (2 years) |
| **Abdominal Aorta** | 6 | Adult (2 years) |
| **Pulmonary Artery** | 6 | Adult (2 years) |

### RNA-seq

The RNA-seq analysis we present in this manuscript is based on a subset of data, from seven cardiovascular and one skeletal muscle tissue (Table 1), from our high resolution atlas of gene expression for domestic sheep (Clark et al. 2017). These tissues were collected from three male and three female adult Texel x Scottish Blackface sheep at 2 years of age (n=6 in total). RNA extraction, RNA-seq library preparation, sequencing and bioinformatic analysis of the RNA-seq data were described in Clark et al 2017. After quality control a small number of samples, despite multiple extraction attempts, were of insufficient quality for RNA-Seq, and as such a small proportion of the tissues have less than six biological replicates (Table 1).

RNA-seq libraries were generated by Edinburgh Genomics (Edinburgh, UK). All libraries were 125bp paired end stranded, sequenced at a depth of >25 million reads per sample, and prepared using the Ilumina TruSeq mRNA library preparation protocol (Ilumina; Part: 15031047 Revision E). The only exceptions were the left ventricle samples that were prepared using the Ilumina TruSeq total RNA library preparation protocol (Ilumina; Part: 15031048, Revision E) and sequenced at a depth of >100 million reads per sample. Supplementary Table 2 includes the details of the cardiovascular tissues included in the sheep atlas dataset and analysed further here.

The raw RNA-seq data are deposited in the European Nucleotide Archive (ENA) under study accession number PRJEB19199 (http://www.ebi.ac.uk/ena/data/view/PRJEB19199). For each tissue a set of expression estimates (averaged across biological replicates from n=6 adult sheep), as transcripts per million (TPM), were obtained using the transcript quantification tool Kallisto v0.43.0 (Bray et al. 2016), as described in Clark et al. 2017. It was necessary to normalise these estimates according to the methods described in (Bush et al. 2017) to account for the two different library types (ribo-depleted total RNA (left ventricle) and poly-A selected mRNA (all other tissues)). The gene expression estimates for the sheep gene expression atlas dataset are publicly available on BioGPS (http://biogps.org/dataset/BDS_00015/sheep-atlas/), and we have included the gene expression estimates for the subset of tissues re-analysed here as Supplemental Dataset 1.

### Visualisation and analysis of the gene expression estimates

Expression estimates from Kallisto for each gene were analysed using the network visualisation tool, Graphia Professional (Kajeka Ltd, Edinburgh, UK; https://kajeka.com/graphia/) (Livigni et al. 2018). Briefly, similarities between individual gene expression profiles, averaged across N=6 biological replicates (3 male and 3 female adult sheep) where possible for each tissue, were determined by the calculation of a Pearson correlation matrix for both sample-to-sample and gene-to-gene comparisons. During this process the dataset was filtered to remove relationships where the Pearson correlation coefficient (which is the statistical measure of the strength of a linear relationship between paired data) was below a threshold of r ≥ 0.91 and r ≥ 0.99, respectively. The Markov clustering algorithm (MCL) was applied at the default inflation value (to determine cluster granularity) of 2.2 (van Dongen & Abreu-Goodger, 2012) to identify groups of transcripts with closely related expression patterns. Clusters were numbered in order of decreasing cluster size. The online Database for Annotation, Visualization and Integrated Discovery (DAVID) Functional Annotation tool (https://david.ncifcrf.gov/) was used for Gene Ontology (GO) analysis.

### Statistical analyses

Statistical analyses were performed using Minitab 17 (Coventry, UK). The Kolmogorov-Smirnov normality test was performed to check whether experimental data were normally distributed. In this study, one-way analysis of variance (ANOVA) using a general linear model incorporating Fisher’s least significant difference (LSD) method was used for pairwise comparisons. Gene expression data in this study are expressed as mean ± standard deviation (SD), and p-value <0.05 was considered significant. Dotplots were made in R v3.2.2 (https://www.r-project.org/), using the R package ‘ggplot2’ with error bars showing mean ± SD. Individual data points are also included in the dotplots.

## Results

To explore the gene expression differences in the whole cardiovascular system, we used RT-qPCR to analyse selected genes involved in extracellular matrix (ECM) composition and in maintenance of calcium homeostasis, in a range of samples from pre- and post-natal developmental stages of the sheep. Genes which are the focus of studies in our group because they are associated with cardiovascular pathology and/or ectopic calcification were examined. The genes and the diseases associated with them are listed in Supplementary Table 3. A summary of the results for the developmental stages is presented in Table 2 and the full results with significance levels can be seen in Supplementary Figures 1-6. Results for different tissues in the adult sheep are summarised in Figure 1 and full results with significance levels can be found in Supplementary Figures 7-9.

**Figure 1:**
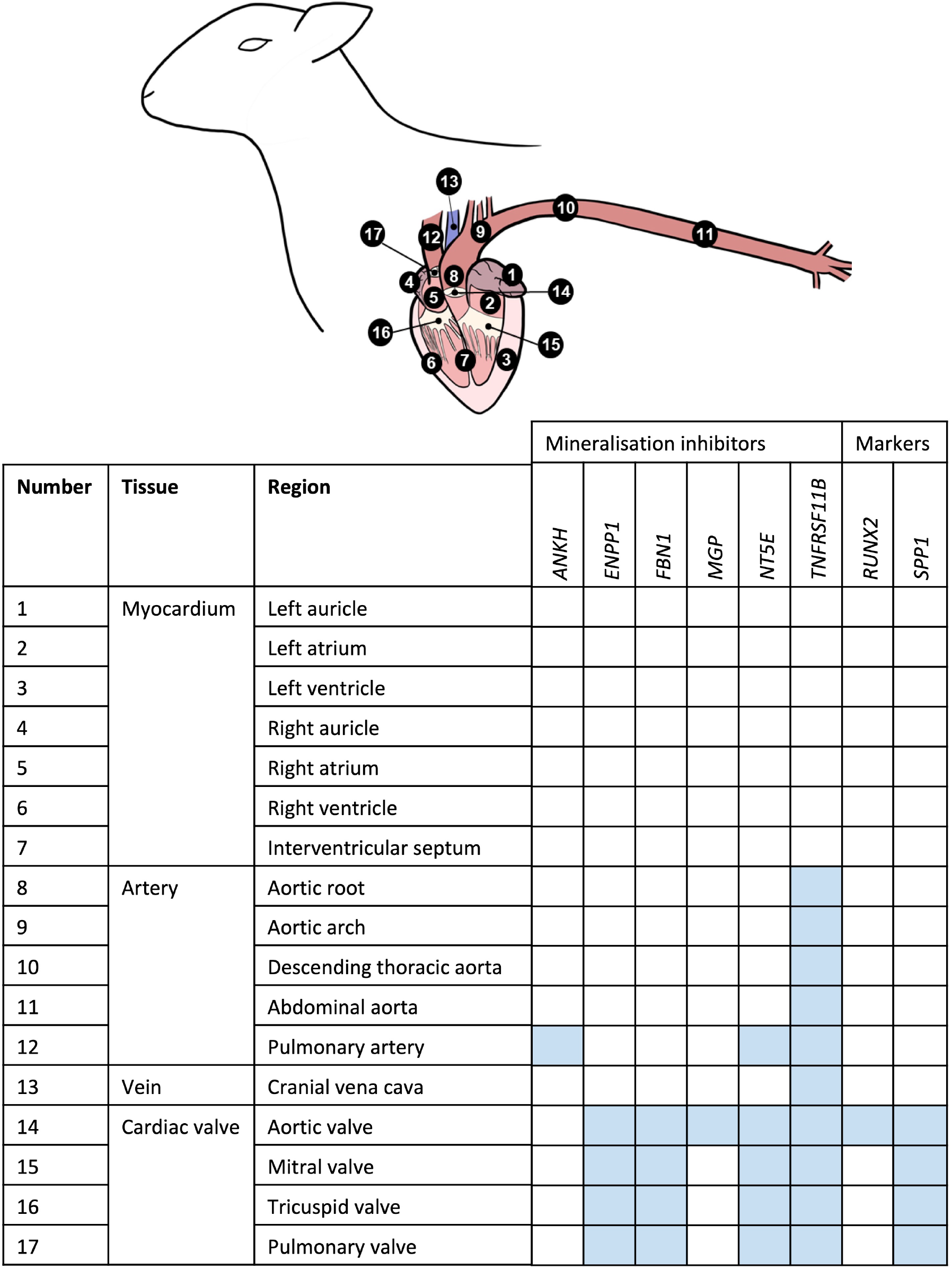
Summary of mRNA expression profiles of key vascular calcification genes in the sheep cardiovascular system. Blue blocks indicate where genes were found to be most highly expressed in this study. AV = atrioventricular.

**Table 2.** Summary of expression profiles of ECM and calcification genes during pre- to post-natal development in the sheep cardiovascular system. Colour key for myocardial tissue only (no foetal samples were available for arterial tissues): Red - Overall changed expression from foetal to adult; Green - Changed expression from pre- to post-natal stages; Blue - Changed expression during post-natal stages.

| Tissue | Genes that increased in expression through development | Genes that decreased in expression through development | Gene(s) with highest expression through development | Gene(s) with lowest expression through development |
| --- | --- | --- | --- | --- |
| *Myocardium* |  |  |  |  |
| Left ventricle | *ANKH* | *COL1A1* | *MMP2* | *TNFRSF11B* |
|  | *RUNX2* | *FBN2* |  |  |
|  |  | *MMP2* |  |  |
|  |  | *SPP1* |  |  |
|  |  | *FBN1* |  |  |
|  |  | *BGN* |  |  |
|  |  | *TIMP1* |  |  |
|  |  | *TNFRSF11B* |  |  |
| Interventricular septum | *ANKH* | *BGN* | *BGN* | *TNFRSF11B* |
|  | *MGP* | *FBN2* | *FBN1* |  |
|  |  | *SPP1* | *MGP* |  |
|  |  | *KCNK3* |  |  |
|  |  | *FBN1* |  |  |
|  |  | *COL1A1* |  |  |
| *Arteries* |  |  |  |  |
| Pulmonary artery | *TIMP1* | *FBN2* | *MGP* | *RUNX2* |
|  | *ENPP1* | *SPP1* |  |  |
|  | *RUNX2* |  |  |  |
| Aortic root | - | *COL1A1* | *MGP* | *RUNX2* |
|  |  | *BGN* |  | *SPP1* |
|  |  | *FBN1* |  | *ANKH* |
|  |  | *FBN2* |  |  |
|  |  | *MMP2* |  |  |
|  |  | *ENPP1* |  |  |
|  |  | *SPP1* |  |  |
|  |  | *RUNX2* |  |  |
|  |  | *KCNK3* |  |  |
| Aortic arch | *TNFRSF11B* | *FBN2* | *MGP* | *RUNX2* |
|  | *MGP* | *SPP1* |  |  |
|  | *RUNX2* |  |  |  |
| Abdominal aorta | *COL1A1* | *FBN2* | *MMP2* | *RUNX2* |
|  | *FBN1* | *SPP1* | *MGP* |  |
|  | *TIMP1* | *RUNX2* | *BGN* |  |
|  | *ANKH* |  |  |  |
|  | *TNFRSF11B* |  |  |  |

### Myocardial tissue gene expression during development

In the left ventricle free wall, the ECM protein-encoding genes showed significant decreases in relative expression during development from 100 days gestation to 2 years of age, although the timing varied, with *BGN* and *TIMP1* declining before birth while *COL1A1, MMP2* and *FBN2* decreased after birth. *FBN1* dropped rapidly before birth but then increased at 8 weeks (Table 2; Supplementary Figure 1). The expression of *SPP1* (also known as *OPN*), which promotes calcification, expression decreased overall with age while *RUNX2*, which is also involved in mineralisation increased during development. The mineralisation inhibitor *ANKH* showed increases in relative mRNA expression, while *TNFRSF11B (*also known as *OPG* and thought to restrict calcificaiton) showed a significant decrease in its expression levels after birth. Overall, the expression levels of *RUNX2* and *TNFRSF11B* were low compared to the other tested genes and *MMP2, BGN* and *MGP* were the highest compared to the other genes in the left ventricle (Table 2; Supplementary Figure 1).

In contrast, in the interventricular septum, the ECM genes *COL1A1, BGN* and *MMP2* were largely unchanged during development while *FBN1* and *FBN2* exhibited decreases in mRNA expression as development progressed (Table 2; Supplementary Figure 2). In general, *SPP1* expression decreased with age in the interventricular septum although higher expression was observed in the 1 week old lambs compared to newborn lambs (p<0.05). As in the left ventricle, the expression of *ANKH* showed an increasing trend with age, but with a significant increase between the foetal and newborn lamb samples (p<0.05). Overall, the expression of *MGP* (mineralisation inhibitor) did not change, although the 1 week old lambs showed statistically significant higher expression compared to the foetal lambs (p<0.05). Genes that were most highly expressed in the interventricular septum were *BGN, MGP* and *FBN1*, whereas *TNFRSF11B* showed the lowest levels of expression within this tissue (Table 2).

### Arterial tissue expression during development

It was not possible to obtain samples from the foetal animals for the arteries, but we examined gene expression changes from newborn to adult. In the pulmonary artery, *FBN2* expression decreased and *TIMP1* expression increased between birth and 2 years of age (Table 2; Supplementary Figure 3). Expression of both *RUNX2* and *ENPP1* (likely to have opposing effects on mineralisation) was significantly higher in the 2-year old sheep (p<0.01; Table 2; Supplementary Figure 3). *SPP1* was significantly lower in the 2-year old sheep compared to both newborn and 8-week old lambs (p<0.01). In the pulmonary artery, *MGP* was the most highly expressed of the genes tested, followed by *BGN, MMP2, ENPP1* and *COL1A1*, with *RUNX2* as the gene with the lowest expression levels (Table 2).

In the aortic root, all the significant changes involved a decrease from young lambs to 2-year-old adults. Both the ECM protein genes (*COL1A1, BGN, MMP2, FBN1* and *FBN2*) and the genes with opposing effects on mineralisation (*ENPP1* and *SPP1)* showed decreases in their expression (Table 2), and were significantly lower at 2 years of age compared to (Supplementary Figure 4). The ion channel gene *KCNK3* also showed significantly lower expression in the 2-year old sheep compared to newborn and 1 week old lambs (p<0.05). Overall, in the aortic root, *MGP* followed by *BGN, MMP2* and *FBN1* showed the highest levels of expression, whereas the lowest levels of expression were observed for *RUNX2* and *SPP1* and *ANKH* in some adult samples (Table 2).

In the aortic arch many of the selected genes were not significantly changed through development (Table 2; Supplementary Figure 5) *FBN2* expression decreased in 2-year old sheep compared to newborn and 1 week old lambs (p<0.01). *SPP1* expression was also significantly lower in adult sheep compared to newborn and 1 week old lambs (p<0.05). In contrast the mineralisation inhibitor genes *TNFRSF11B* and *MGP* increased with age as did *RUNX2.* Within the aortic arch, the highest levels of expression were seen in *MGP, BGN* and *ENPP1*, and the lowest in *RUNX2* and *TNFRSF11B* (newborn lambs) and *FBN2* and *SPP1* (2-year old adults) (Table 2).

In the abdominal aorta, expression of the mRNAs of ECM protein-encoding genes *COL1A1* and *FBN1* peaked in 8-week old lambs (Supplementary Figure 6) while *FBN2* expression decreased from 8 weeks of age to 2 years of age (p<0.01). *TIMP1* expression was found to increase with age. For the key calcification genes, *SPP1* showed a reduction in its expression from 8-week old lambs to 2-year old sheep (p<0.05), and *RUNX2* was decreased from newborn to 8-week old lambs (p<0.05; Supplementary Figure 6). *ANKH* and *TNFRSF11B* expression was significantly increased in the 8-week old and 2-year old sheep compared to newborn lambs (p<0.05). In the abdominal aorta, the highest levels of expression were observed for *MMP2, MGP, BGN* and *FBN1*, and the lowest overall for *RUNX2* (Table 2).

### Vascular calcification (VC) inhibitors expressed in the healthy adult cardiovascular system

Using RT-qPCR the gene profiles of various key calcification inhibitors were investigated in different cardiovascular regions, in the six adult sheep. Figure 1 summarises where these genes were highly expressed in the cardiovascular tissues. Overall, the key VC genes tended to be more highly expressed in the cardiac valves than the other tissues; expression in the myocardium was lowest (Figure 1).

Of the genes examined, *FBN1 and TNFRSF11B* showed the greatest variation across the cardiovascular system. *FBN1* was most highly expressed in the valves, which had significantly higher expression than the myocardium and vena cava tissues (p<0.01; Figure 3). Overall, *FBN1* expression was approximately 4-fold lower in the aortic samples than the cardiac valves (p<0.05), There was no significant difference between the aortic arch compared to the left AV valve, the abdominal aorta compared to the left AV (mitral), right AV (tricuspid) and pulmonary valves, and the pulmonary artery compared to the left AV valve (Figure 2). Expression of *FBN1* in the arteries was in general higher (4 fold) than in the myocardium and cranial vena cava (p<0.05). *TNFRSF11B* expression was highest in the arteries and cardiac valves compared to the myocardium and cranial vena cava (Figure 2). The levels of *TNFRSF11B* expression in the aortic samples, pulmonary artery, the aortic valve and left AV valve were significantly higher compared to the myocardium and cranial vena cava (p<0.05; Figure 2). The expression of *TNFRSF11B* was generally very low in the myocardium (approximately 1000-fold lower than in the arteries) (Figure 2).

**Figure 2:**
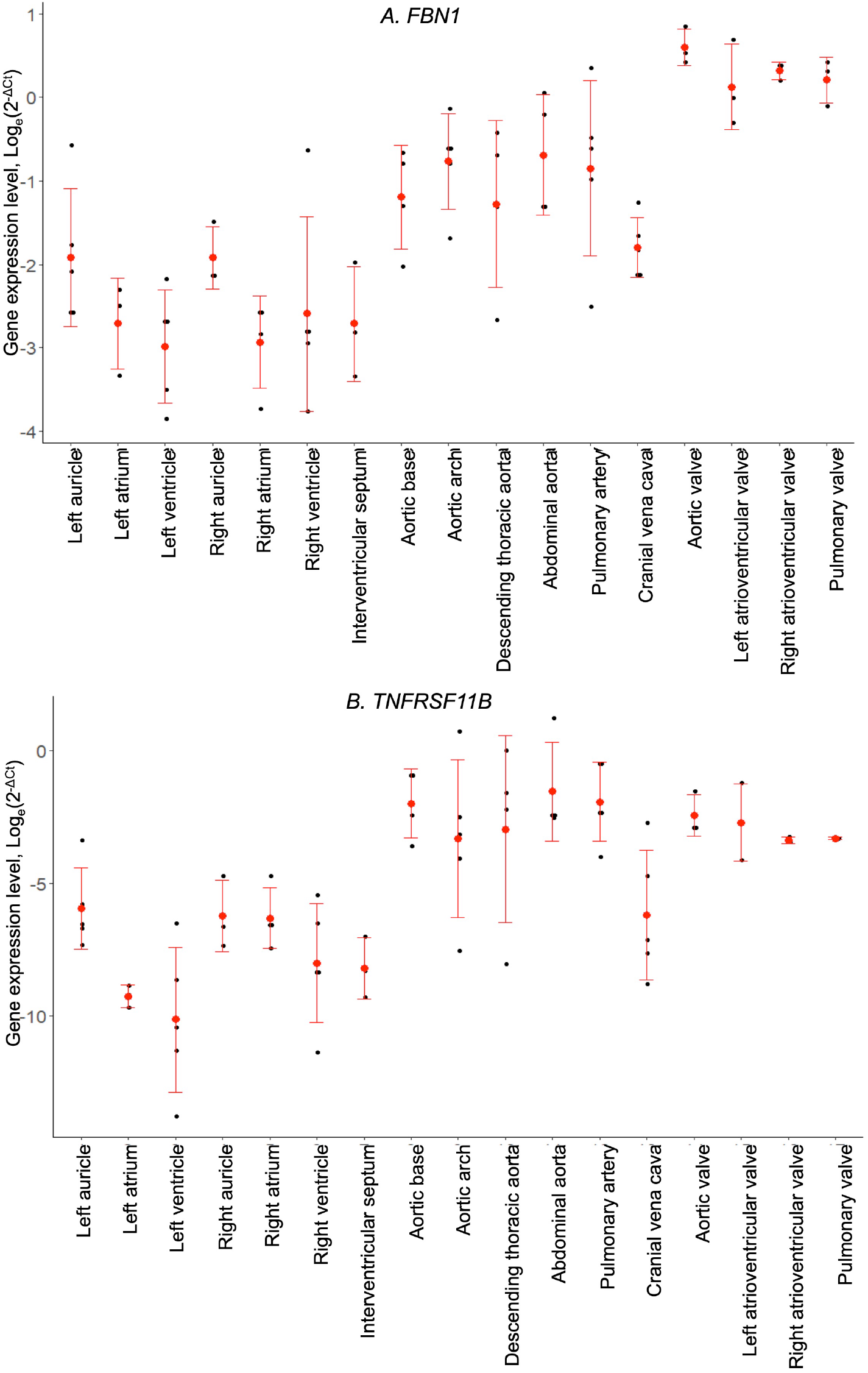
mRNA expression levels for individual animals, determined by RT-qPCR. (A) fibrillin-1 (*FBN1*) and (B) osteoprotegerin (*TNFRSF11B*). Gene expression levels were normalised to the geomean of *GAPDH* and *YWHAZ*. Dot plots show individual data points (black dot), the mean expression for each tissue (red dot) and standard deviation (red error bars).

**Figure 3.**
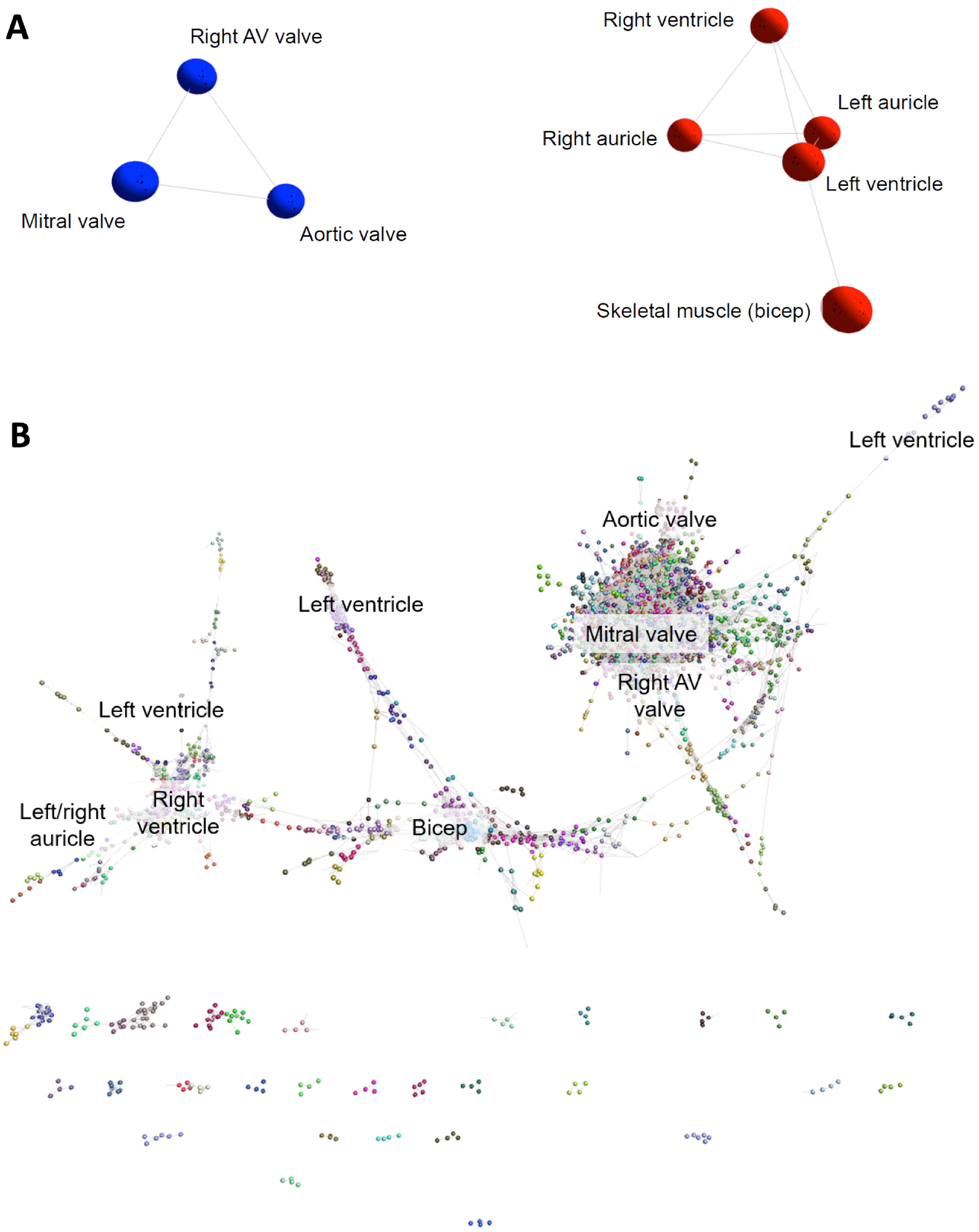
Cardiovascular gene co-expression networks. (A) Sample-to-sample analysis. Network layout of 8 tissue types revealed two distinct components: the first containing cardiac valve samples (blue), and the second with myocardium/skeletal muscle tissues (red). Pearson correlation co-efficient r ≥ 0.91. (B) Gene-to-gene analysis. Gene clusters are distinguished by colour, and the labels indicate clusters of genes most highly expressed in the different tissues. With a Pearson correlation coefficient of r ≥ 0.99, the gene-to-gene network was comprised of 11,341 nodes (genes) with 938,652 connections. Markov clustering algorithm (MCL) clustering of the graph (inflation value 2.2) resulted in 555 clusters (containing >3 genes).

Of the other VC genes investigated, *MGP* showed the highest levels of expression compared to the other tested genes with expression being similar in all tissues, although the aortic valve showed higher expression than the aortic arch, the cranial vena cava and the myocardial tissues (p<0.05; Supplementary Figure 7). The expression of *ANKH* was similar in all tested tissues, but the pulmonary artery showed significantly higher expression levels than the right atrium and left auricle (p<0.05; Supplementary Figure 7). *NT5E* expression was found to be higher in the cardiac valves and the pulmonary artery compared to the other tissues included in this study (p<0.05; Supplementary Figure 8). Some arterial tissues exhibited higher expression than the myocardial samples, including the aortic root, aortic arch and abdominal aorta compared to the right atrium, left ventricle and interventricular septum (p<0.05; Supplementary Figure 8). The expression of *ENPP1* was significantly higher in the cardiac valves and aortic arch compared to the myocardium and vena cava (p<0.05; Supplementary Figure 9). The remaining aortic samples showed intermediate expression between the valves and myocardium (Supplementary Figure 9). Similarly the expression of *SPP1* was found to be higher in the cardiac valves compared to the myocardium and the aortic root (p<0.05; Supplementary Figure 9). Other than in the cardiac valves, *SPP1* expression was generally very low in the tested cardiovascular tissues, reaching levels similar to that of the bone marker *RUNX2* (Supplementary Figure 9). The expression of *RUNX2* was low in all tested samples. However, the aortic valve showed higher *RUNX2* expression compared to the myocardium, the cranial vena cava and aortic root (p<0.05; Supplementary Figure 8).

### Gene expression profiles reflect anatomical structure

In addition to analysis of selected ECM and VC genes across a wide range of tissues and developmental stages, we took advantage of the RNA-seq data from the sheep atlas project (Clark et al. 2017) to explore more broadly the transcriptome of the heart and cardiac valves. The 3D network visualisation for sample-to-sample analysis is similar to a principal components analysis and grouped the tissue samples (averaged across biological replicates from up to 6 adult sheep; Supplemental Dataset 1) together based on highly similar expression profiles. The resultant graph contained all 8 nodes (tissue samples) that were connected by 10 edges (connections between nodes at a correlation coefficient of ≥0.91; Figure 3A). Two distinct elements were identified: a group containing the five myocardium/skeletal muscle samples and a cardiac valve group (Figure 3A). The network indicated that there were close similarities in the overall expression profiles of genes in skeletal muscle and heart muscle, which were distinct from the cardiac valve tissues. Similar grouping of skeletal muscle and heart muscle tissues was previously observed by (Lukk et al. 2010).

### Tissue-specific expression clusters in the cardiovascular system

Network-based gene-to-gene analysis of Supplemental Dataset 1 grouped genes according to their expression pattern across the muscle and valve samples, producing a gene co-expression network (GCN) (Gaiteri et al. 2014). A high correlation coefficient of r ≥ 0.99 was necessary to discriminate expression npatterns due to the similarity of expression in the relatively small set of samples being analysed. The resultant graph (Figure 3B) included 11,341 nodes (genes) with 938,652 edges (correlations at r ≥ 0.99 between them). MCL (inflation = 2.2) clustering of the graph resulted in 555 clusters containing >3 genes. Details of the genes included in the 40 largest clusters are included in Supplemental Dataset 2. For most of the clusters, expression was highest in the three valve samples. Some of these (where the difference in average expression was less than 4-fold) contained housekeeping genes and were considered ubiquitous, while others (where the difference was 10-fold or more) were considered to be valve specific. Several clusters contained genes that were high in skeletal muscle only and some were high in the myocardium samples only. There was no cluster of genes that were high in the ventricles alone, or in a single myocardium sample, but a small number of clusters contained genes that were high in the auricles. Interestingly there were some clusters of genes with higher expression in the skeletal muscle and heart valves than the myocardium or with higher expression in the skeletal muscle and left ventricle. There were also some differences between the different valves. Expression profiles of a subset of ECM, VC and heart function genes are shown in Supplementary Figure 10..

Cluster 1 contained 3543 genes, with 529 of those unannotated (Figure 4A). Genes in this cluster showed approximately a 3-fold greater expression in the cardiac valves than in the other tissues. A wide variety of genes was included in this cluster. The cluster contained genes enriched for GO terms associated with cell structure, such as *COL1A1* (encoding collagen type I alpha 1) and *COL3A1* (collagen type III alpha 1), *MMP*2, -*9*, -*19*, -*20* and -*28* (matrix metalloproteinases), *FBN2* (fibrillin-2) and *TIMP1* (tissue inhibitor of matrix metalloproteinases 1) (Table 3). There were also multiple genes expressed specifically by macrophages in sheep (Clark et al. 2017) and other species (Fantom Consortium et al. 2014; Summers et al. 2020), including *CSF1R, AIF1* and *SPI1*, *CSF2RA/B* and a number of interleukin and interferon responsive genes. In addition, members of the smoothened signalling pathway (*SMO, GLI1-3*) involved in cilium formation and function, genes associated with RNA transcription and processing (for example POLR genes) and some genes associated with cell proliferation (for example centromere protein genes) were in this cluster. Cluster 1 also contained genes associated with TGF beta signalling, including *TGFB3, TGFBI, TGFBR2* and *TGFBR3.*

**Figure 4.**
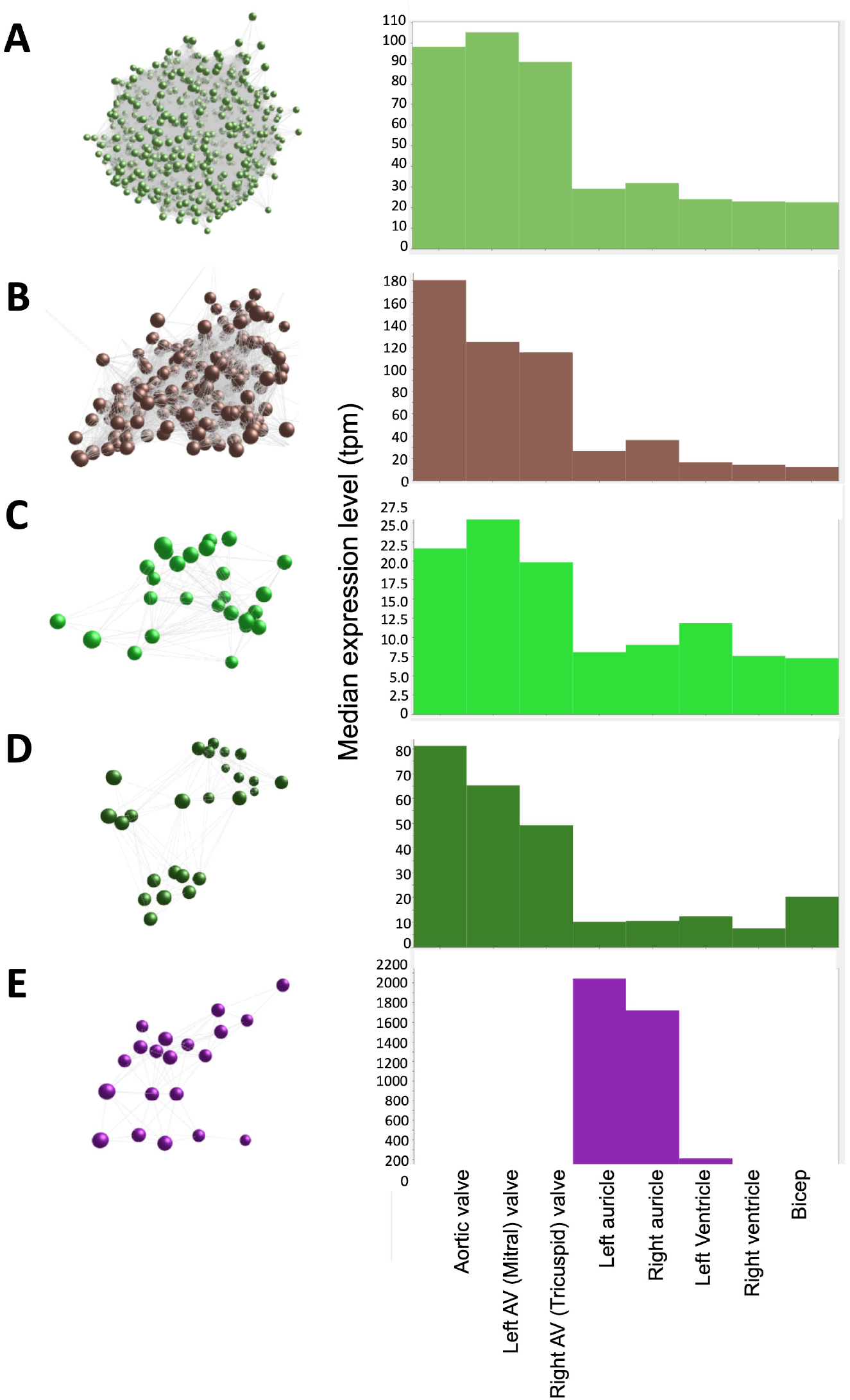
Average expression of genes within clusters. X axis shows the samples; Y axis shows the average expression for the cluster in tpm. Spheres (nodes) in co-expression clusters (left) denote individual genes; lines represent connections between genes. The histogram (right) shows median expression levels in transcripts per million (TPM) in the different tissues. AV = atrioventricular. (A) Cluster 1, the largest cluster, contained 3543 genes, with 529 unannotated genes. A co-expression cluster highly expressed in the sheep cardiac valves compared to the myocardium and bicep. (B) Cluster 3 contained 192 genes, with 40 unannotated genes. A co-expression cluster highly expressed in the sheep cardiac valves, particularly in the aortic valves, compared to the myocardium and bicep. (C) Cluster 22 contained 27 genes, with 5 unannotated genes. A co-expression cluster highly expressed in the sheep cardiac valves compared to the myocardium and bicep. (D) Cluster 24 contained 25 genes, with 3 unannotated genes. A co-expression cluster highly expressed in the sheep cardiac valves compared to the myocardium and bicep. (E) Cluster 36 contained 20 genes, with 5 unannotated genes. A co-expression cluster highly expressed in the sheep auricles compared to the cardiac valves, the ventricles and the bicep.

**Table 3.** Summary of 5 clusters from the sheep cardiovascular transcriptome dataset. Gene Ontology (GO) term analysis was performed using DAVID Functional Annotation tool (https://david.ncifcrf.gov/). BP = Biological Processes; CC: Cellular component; MF: Molecular Function. Full gene lists and cardiovascular expression data are presented in Supplemental Dataset 1. *Up to 3000 genes maximum can be input into DAVID, thus two runs were performed with the ^a^top 3000 genes and then ^b^bottom 3000 genes. EASE score = modified Fisher Exact. > indicates decreasing expression.

| Cluster | Number of genes | Number of un-annotated genes | Expression profile description | Functional class | GO term | EASE score (p-value) | EASE score (p-value; Benjamini corrected) | Genes included (Gene symbols) |
| --- | --- | --- | --- | --- | --- | --- | --- | --- |
| 1 | 3543 | 529 | Cardiac valves | Various, Housekeeping | * (BP) mRNA 3'-end processing; RNA export from nucleus | <sup>a,b</sup>8.1E-6;<br><sup>a,b</sup>1.5E-4 | <sup>a,b</sup>0.0064;<br><sup>a</sup>0.063,<br><sup>b</sup>0.055 | COL1A1, COL3A1, MMP2, TIMP1 |
| 3 | 192 | 42 | Cardiac valves (highest in aortic valve) | ECM organisation, bone development | (BP) extracellular matrix organization; skeletal system development; osteoblast differentiation | 6.02E-4;<br>0.00336;<br>1.12E-5 | 0.124;<br>0.31;<br>0.0099 | BGN, COL1A2, SPARC, BGLAP, COL1A2, GDF10, BMP4, BGLAP, SNAI1-2, SOX8 |
| 22 | 27 | 5 | Cardiac valves | Housekeeping, Immune | (BP) immune response<br>(MF) integral component of membrane | 0.067;<br>0.061 | 0.999;<br>0.928 | CCR6, ENPP1, NFIL3, CD47, ENPP1, IL6ST, SMAD2 |
| 24 | 25 | 3 | Cardiac valves (Aortic valve > Mitral valve > Tricuspid valve) | ECM | (BP) positive regulation of transcription from RNA polymerase II promoter; transcription from RNA polymerase II promoter<br>(CC) extracellular matrix | 0.108;<br>0.123;<br>0.045 | 0.999;<br>0.999;<br>0.943 | MEOX1, TRPS1, RBPJ, NLRP3, FMOD, FBN1, COL6A1 |
| 36 | 20 | 5 | Auricles (Left > Right) | Muscle contraction | (BP) potassium ion transport;<br>(MF) calcium ion binding;<br>(CC) extracellular space | 0.002;<br>0.06;<br>0.221 | 0.16;<br>0.921;<br>0.999 | KCNQ3, KCNJ3, KCNK3, MYL7, PAM, MYL4, DKK3, NPPA, PAM |

Cluster 3 (Figure 4B) contained 192 genes with highest expression in the cardiac valves, particularly the aortic valve, at approximately 4.5 to 18-fold higher than the myocardial and skeletal muscle tissues. This cluster was enriched for GO terms associated with extracellular matrix organisation, skeletal system development, osteoblast differentiation and cartilage morphogenesis. Genes in this cluster included those encoding bone morphogenetic protein 4 (*BMP4*), collagen type I alpha 2 (*COL1A2*) and bone gamma-carboxyglutamate protein (*BGLAP*, also known as osteocalcin) (Table 3). Other valve specific genes in this cluster included *MYH10* (myosin heavy chain 10), Potassium channel gene *KCNU1* and *BGN* (ECM protein biglycan) which was 15- to 20-fold higher in the valves. Genes encoding various transcription factors were also contained within this cluster, including FOS like 2, AP-1 transcription factor subunit (*FOSL2*), Twist basic helix-loop-helix transcription factor 2 (*TWIST2*), Snail family transcriptional repressor 1 and 2 (*SNAI1 and -2*) and Sry homeobox 8 (*SOX8*). In cluster 3, 42/192 genes were unannotated and not included in the input into DAVID.

Like cluster 1, the 27 genes in cluster 22 (Figure 4C) also showed relatively ubiquitous expression, with smaller differential between the cardiac valves and the myocardium and skeletal muscle than those in cluster 1 (approximately 2 to 3-fold difference) (Figure 4C). Genes in cluster 22 included *ENPP1* (ectonucleotide pyrophosphate/phosphodiesterase 1), *ADAMTS6* (ADAM metallopeptidase with thrombospondin type 1 motif 6) and *SMAD2* (SMAD family member 2). Some of the genes in this cluster are annotated as immune-related, including C-C motif chemokine receptor 6 (*CCR6*), interleukin6 signal transducer (*IL6ST*) and nuclear factor, interleukin 3 (*NFIL3*) (Table 3) but these genes are less specific to hematopoietic cells than those in Cluster 1 (https://www.bioGPS.org/sheepatlas; Clark et al. 2017).

The 25 genes in cluster 24 (Figure 4D) were also more highly expressed in the cardiac valves, at up to 16-fold higher than the myocardium, and approximately 4 to 5-fold higher than the bicep (representing skeletal muscle). Genes encoding proteins involved in transcriptional regulation were included in this cluster, including mesenchyme homeobox 1 (*MEOX1*), the GATA-regulated gene repressor transcriptional repressor GATA Binding 1 (*TRPS1*) and recombination signal binding protein for immunoglobulin kappa J region (*RBPJ*) (Table 3). ECM protein encoding genes were also included, such as *FBN1* (fibrillin-1), *FMOD* (fibromodulin), and *COL6A1* (collagen type VI alpha 1) as well as the macrophage specific gene NLR family pyrin domain containing 3 (*NLRP3*) (Table 3).

The 20 genes in cluster 36 (Figure 4E) showed high expression in the auricles, at up to 2,000-fold higher than the other tissues. Genes included in this cluster were involved in muscle contraction through potassium ion transport ion transport (Table 3) e.g. potassium voltage-gated channel subfamily Q member 3 (*KCNQ3*), subfamily J member 3 (*KCNJ3*) and subfamily K member 3 (*KCNK3*), as well as peptidylglycine alpha-amidating monooxygenase (*PAM*) and myosin light chains 4 and 7 (*MYL4 and -7*) (Table 3). Other genes to note include natriuretic peptide A (*NPPA*) and Dickkopf WNT signalling pathway inhibitor 3 (*DKK3*) (Table 3). Examination of the wider sheep gene expression atlas (https://www.biogps.org/sheepatlas) showed that many of the genes in this cluster were also highly expressed in brain regions, consistent with the function of many as ion transport channels.

A number of clusters contained genes that were high in both bicep (representing skeletal muscle) and the heart regions and 2- to 3-fold lower in the valves. As might be expected, these clusters were enriched for genes involved in mitochondrial function, reflecting the energy requirements of muscle tissue. In contrast, Cluster 4 (164 nodes) expression was high only in bicep and contained genes specific to skeletal muscle such as a range of troponin and myosin genes, the ryanodine receptor 1 gene (*RYR1*) and sodium, potassium and calcium ion channels genes. Other bicep-specific clusters included Clusters 23 (25 genes) and 28 (24 genes). These three clusters were comprised of genes encoding proteins for zinc-dependent proteases e.g. *ADAMTS20* (ADAM metallopeptidase with thrombospondin type 1 motif 20) and genes involved in actin biding and motor activity e.g. *MYH15* (myosin heavy chain 15). A small proportion of clusters exhibited expression patterns that were specific to skeletal muscle (bicep) and left ventricle. The largest of these clusters was cluster 13 (48 genes), which included genes encoding proteins involved in hydrolysis of extracellular nucleotides e.g. ectonucleotide pyrophosphatase/phosphodiesterase 3 (*ENPP3*) and transcription factors e.g. caudal type homeobox 1 (*CDX1*), solute carriers e.g. (*SLC16A4* and *SLC26A3*). Several smaller clusters, 66 (15 genes), 83 (13 genes) and 90 (13 genes), exhibited left ventricle specific expression profiles and were comprised of genes with a similar function to those within cluster 13.

In summary, the cardiac valves showed more consistent expression of ECM genes, while both valves and the muscle samples had detectable levels of many transcripts associated with resident macrophage populations. The transcriptome of cardiac muscle was similar to skeletal muscle in this analysis, although some tissue specific gene expression could be seen. For example, the potassium channel genes *KCNJ3, KCNK3* and *KCNQ3*, the myosin light chain genes *MYL4* and *MYL7* and a natriuretic peptide gene, *NPPA* were auricle-specific (Cluster 36).

### Functional annotation of unannotated genes

Each cluster included a number of genes, which had no informative gene name. While many of these genes had low expression, some were highly expressed. For example, *ENSOARG00000020353* had a maximum of nearly 6,000 TPM in aortic valve. It is described as a novel gene with a 24 amino acid match to parathymosin (*PTMS*) in the bovine. According to the ‘guilt by association’ principle (Oliver et al. 2000) since this gene was found within a cluster of genes with high expression in the cardiac valve samples it may well have a similar function to other annotated genes that are co-expressed in Cluster Other examples include *ENSOARG00000005484* in Cluster 36, which was expressed at around 18 TPM in left and right auricle. This gene has some homology to *FAM155A* and *FAM155B* (*Tmem28* in mouse), probably a transmembrane calcium ion transporter (UniProtKB B1AL88 and O75949 respectively). Further exploration of the 923 ENSOARG genes that were included in the cluster analysis should allow attribution of putative functions based on their presence in a cluster of genes of known function.

## Discussion

The maintenance of a healthy cardiovascular system requires expression of genes that contribute to essential biological activities and repression of those that are associated with functions likely to be detrimental to cardiovascular homeostasis. As cardiovascular disease (CVD) is of major clinical importance, understanding the roles of genes in co-expression networks and their associated molecular pathways will be useful in understanding their dysregulation in pathological events. Detailed analysis of gene expression of tissues and cell types in the cardiovascular system provides a powerful resource for investigation of healthy cardiovascular system function (reviewed in Wirke et al. 2018). However, analysis of the cardiovascular system in humans is often jeopardised due to tissues being only available post-mortem, where frequently the health status of the individual is not known and the quality of the RNA may be poor (Ferreira et al. 2017). A large animal model, where tissues can be collected quickly post mortem from healthy animals, offers the opportunity to perform a detailed characterisation of the mammalian cardiovascular transcriptome. The recently published sheep gene expression atlas (https://www.bioGPS.org/sheepatlas; Clark et al. 2017) allowed us to examine individual components of the cardiovascular system in the sheep, which are similar to humans in their physiology and genetics (reviewed in Hamernik 2019). Additional tissues and developmental stages, from the same animals, but not initially presented as part of the sheep transcriptional atlas, were also examined in this study, using RT-qPCR. The insights from this novel analysis of the sheep cardiovascular system will be valuable in understanding the mechanisms behind many cardiovascular-related diseases and will help to facilitate the development of clinical and therapeutic approaches for the prevention and treatment of cardiovascular diseases.

Our results demonstrate that there is extensive expression of genes encoding proteins involved in formation and maintenance of the extracellular matrix (ECM) in cardiovascular tissues. The ECM is an important provider of structural and biomechanical support, and helps to regulate molecular interactions between growth factors and cell surface receptors (Davis & Summers, 2012; Kim, Turnbull, & Guimond, 2011). The cardiac valves showed higher expression of a range of ECM genes relative to cardiac muscle, both by RNA-seq and by RT-qPCR. This is consistent with continuous remodelling of the ECM in cardiac valves due to the normal functional stresses (mechanical and blood-flow induced shear) that the valve is subjected to. During development, most ECM genes decreased in expression, particularly in the left ventricle, intraventricular septum and aortic root. This probably reflects the completion of organ development, after which the requirement for expression would depend on the turnover of the proteins to maintain ECM homeostasis. However, with ageing, expression and *de novo* synthesis may not be sufficient to balance turnover of the proteins, leading to a loss of structural support in the ECM over time. Different ECM genes were activated at different times during development. For example, two members of the fibrillin family, key components of the ECM (Davis and Summers 2012; Ramirez & Pereira, 1999; Sakai, Keene, & Engvall, 1986) were expressed in the cardiovascular system. The gene encoding FBN2, traditionally regarded as a fetal protein which is involved at the beginning of elastogenesis and early morphogenesis (Zhang et al. 1995), appeared to be activated earlier in cardiovascular development than the gene for FBN1, which has been attributed functions late in morphogenesis and organogenesis (Zhang et al. 1995). Both genes exhibited highest expression in the cardiac valves and decreased in expression during cardiovascular development. Expression of *FBN1* in the aorta supports the role of FBN1 in maintaining the structural integrity of this major artery. In Marfan Syndrome, the dysfunction of FBN1 leads to aortic aneurysms and elastic fibre calcification (Pereira et al. 1999; Bunton et al. 2001). Other ECM genes that decreased with age included *COL1A1* and *BGN.* The ECM proteases known as matrix metallopeptidases (MMPs) and their tissue inhibitors, TIMPs, are important modulators of matrix protein turnover (Elmore, Keister, Franklin, Youkey, & Carey, 1998; Hughes & Jacobs, 2017). It is thought that alterations of the balance between MMPs and TIMPs are critical in the formation of aortic aneurysms and age-associated physiological changes in the cardiovascular system (Meschiari, Ero, Pan, Finkel, & Lindsey, 2017; Rabkin, 2014). In this study, the level of *MMP2* expression was amongst the highest of the genes examined in the abdominal aorta, and *TIMP1* expression was found to increase with age in this tissue. The increase in *TIMP1* expression may help prevent the development of abdominal aortic aneurysms in a healthy animal by inhibiting degradation of ECM structural proteins. MMPs and TIMPs may also be crucial in myocardial function, where increases in their levels have been found to correlate with age in human and mouse (Meschiari et al. 2017). Furthermore, these ECM regulators have also been reported to be important in the remodelling process in the left ventricle after experimentally induced myocardial infarction in mice, where the local endogenous control of MMPs by TIMP1 was suggested to be important for the ECM structure, as well as myocardial function and myocyte growth (Creemers et al. 2003). Additional studies on the expression of other MMPs and TIMPs may be useful to determine their involvement in the development of CVD.

We detected transcripts associated with macrophages in all samples, notably enriched in the valves. The homeostatic functions of resident macrophages in arterial and cardiac tissue have been widely studied (Swirski et al. 2016; Lim et al. 2018). The presence of resident macrophages in human and mouse valve tissue has also been recognised previously (Sraeyes et al. 2018). In the mouse, heterogeneous resident valve macrophage populations are established in the postnatal period and the population is expanded by monocyte recruitment in a model of myxomatous disease (Hulin et al. 2018). Damaged cardiac valves are prone to life-threatening infectious and non-infectious endocarditis (Yang and Frazee 2018), which is common in elderly humans, and ongoing surveillance and repair are necessary to prevent pathological outcomes. The sheep is an ideal animal to investigate the ageing heart further, since the sheep life span is around 10 years (http://www.sheep101.info) and elderly animals can be obtained from commercial sources at the end their productive life, rather than needing to be aged for the investigation.

A striking finding of this analysis was the expression of genes associated with bone formation and VC in the cardiovascular system of healthy sheep throughout development. These included both genes encoding proteins that promote bone formation and calcification (such as *SPP1, SPARC, BMP4* and *BGLAP*) and those which suppress mineralisation (such as *ENPP1, ANKH, FBN1, MGP, TNFRSF11B* and *NT5E*). VC can develop in various tissues, although many reports include the aorta and the aortic valve as sites of VC (L. L. Demer & Tintut, 2009; New & Aikawa, 2011). The expression of genes associated with suppression of bone formation would likely be advantageous in preventing VC, but the predisposing factors and pathways that infer the susceptibility of specific tissues to calcification are still unknown. Moreover, differences in the mechanisms behind intimal, median and valvular calcification may exist (Côté et al. 2012; Patel et al. 2017; Qian et al. 2017). Expression of *ENPP1* decreased throughout development. ENPP1 has a role in regulating extracellular nucleotide levels and potentially a dual role in VC (Côté et al. 2012). *ENPP1* may contribute to normal cardiovascular function through the regulation of extracellular ATP concentrations and the generation of the calcification inhibitor PPi (Côté et al. 2012; Nam et al. 2011). Deficiency of *ENPP1* leads to generalised arterial calcification (Mackenzie et al. 2012). *ANKH* mRNA was also increased. *ANKH* transports cytoplasmic PPi out of the cell (Harmey et al. 2004). *ANKH* may provide a protective effect against the development of VC, since patients with VC have been found to have decreased *ANKH* expression (Zhao et al. 2012). MGP is also a calcification inhibitor, possibly via its ability to block BMP signalling (Yao et al. 2010; Zebboudj, Imura, & Bostrom, 2002). Expression of *BMP4* in the cardiac valves in the sheep gene expression atlas dataset analysed here, supports the importance of calcification inhibitors like MGP in preventing the development of calcification, especially in tissues which express genes associated with bone development. The expression of *MGP* was consistently high in all the different ages and tissues investigated and this factor may play a cardioprotective role against the development of calcification. *SPP1* encodes secreted phosphoprotein 1, also known as osteopontin which is associated with bone formation and calcification, and is a constituent of normal elastic fibres in the aorta and skin (Rutsch et al. 2011). SPP1 in the valves is likely to be associated with the resident macrophages, since it was the most highly-expressed transcript in isolated macrophages in the sheep atlas, at least 100-fold higher than in any tissue other than placenta (https://www.bioGPS.org/sheepatlas; Clark et al. 2017). *SPP1* is similarly macrophage-enriched in humans (Fantom Consortium et al. 2014) and pig (Summers et al. 2020). Increased *SPP1* mRNA expression and plasma osteopontin levels have been linked with Cardiac Allograft Vascular Disease (CAVD) (Rajamannan et al. 2003; Yu et al. 2009), whereas it has been reported to have inhibitory effects on arterial calcification (Speer et al. 2002; Wada, McKee, Steitz, & Giachelli, 1999). Examples of its reported roles include bone remodelling, anti-apoptotic signalling and inflammatory regulation (Denhardt, Noda, O’Regan, Pavlin, & Berman, 2001). SPP1 can exist in different states (phosphorylated and glycosylated), and it is thought that these specific forms have distinct functions (Denhardt et al. 2001). There was a decrease in expression of *SPP1* mRNA with age in the sheep cardiovascular tissues. Increased expression of *SPP1* has been implicated in VC development, as well as coronary artery disease and heart failure (Dai et al. 2014; New & Aikawa, 2011; Rosenberg et al. 2008). Our results suggest that *SPP1* is important in the earlier stages of cardiovascular development, whereas higher expression in later life may lead to these adverse clinical outcomes. *ENPP1* was also strongly macrophage-enriched in the wider sheep atlas (https://www.bioGPS.org/sheepatlas; Clark et al. 2017). The expression profiles of *SPP1* and *ENPP1* were very similar suggesting that they contribute to a balance between promotion and suppression of calcification in cardiovascular tissues. *MGP* expression was high compared to the other genes in this study. Although it has been established that *MGP* has a role in the inhibition of VC, its particular role within the cardiovascular system is still unclear. As with SPP1, *MGP* can exist in different states, and the levels of these different states are thought to affect the CVD risk of an individual (Dalmeijer et al. 2013). Elevated dephosphorylated *MGP* (*dpMGP*) has been found in patients with chronic kidney disease (CKD), heart failure, CAVD, aortic stenosis and other CVD events (Mayer et al. 2014; Schurgers, Cranenburg, & Vermeer, 2008; Vassalle & Iervasi, 2014). The locally produced active form of MGP (phosphorylated and gammacarboxylated) has been implicated to have cardioprotective effects (El Asmar, Naoum, & Arbid, 2014; Y. P. Liu et al. 2015; Schurgers et al. 2010) such as through its inhibition of VC, where it has been reported to inhibit BMP signalling (Yao et al. 2010; Zebboudj et al. 2002). In addition, decreased active MGP was found in aortic valvular interstitial cells (VICs) derived from patients with CAVD (Venardos et al. 2015). More studies into the numerous genes that have been implicated in VC are required in order to understand their expression patterns within the cardiovascular system, and to gain additional insights into their physiological functions.

One outcome of this study is the functional annotation of previously novel genes. At present, there are many predicted mammalian protein-coding loci and non-protein-coding genes that are yet to have informative annotation (Oliver 2000; Klomp & Furge, 2012). Protein-coding genes that contribute to common generic and cell-specific cellular processes or pathways generally form co-expression clusters, allowing the inference of the function of a gene (of previously unknown function) using the ‘guilt-by-association’ principle (Oliver 2000; Freeman et al. 2012; Klomp & Furge, 2012). Martherus et al. 2010, for example, used this method effectively to identify heart enriched mitochondrial genes. In our study a number of co-expression clusters were found that distinguished the cardiac valves from heart muscle. The novel (unannotated) genes within the tissue-specific clusters described here potentially have the same functions as other genes in the cluster, which allows for functional annotation of these genes. For example, the gene *ENSOARG00000005484* from Cluster 36 encodes a protein involved in calcium ion transport across membranes, consistent with the other ion channel genes in this cluster. The high level of expression of some of these novel genes suggests that they are an important part of the process of development and differentiation in the cardiovascular system. As such they warrant further investigation using knock out animals or functional validation in relevant cell lines using CRISPR to examine consequences of their dysfunction (as reviewed in Van Kampen and Rooij 2019).

As the RNA-seq analysis we present here was only performed for seven cardiovascular tissues (three valves and the four chambers of the heart), we were not able to define gene expression clusters associated with other cardiovascular tissues, such as the veins, arteries and other regions of the heart. We used RT-qPCR to examine a limited number of genes in the extended cardiovascular system at several developmental stages. Transcriptomic analysis using RNA-seq of a wider sub-set of samples, including more tissue types and developmental stages, would identify specific expression patterns, for example for different parts of the aorta. In addition, we did not cluster the cardiovascular samples with other tissues (other than a representative of skeletal muscle) from the wider sheep gene expression atlas dataset (https://www.bioGPS.org/sheepatlas; Clark et al. 2017).

In summary we have used RNA-seq results from the sheep heart and cardiac valves to further explore the transcriptome of the cardiovascular system in this large animal. These data provide initial insights into tissue-specific expression of key genes, which will be useful in understanding their physiological function in a healthy mammal. This study will support future research into the functions of implicated genes in the development of VC, and increase the utility of the sheep as a model in cardiovascular research. The analysis of further tissues and developmental stages, such as a wider range of prenatal ages and elderly animals would provide further insight into the gene expression patterns of key genes implicated in the progression of important cardiovascular functions or disease with age, and is feasible using the sheep as a model. Here we have built a foundation to explore the transcriptome of the developing and ageing cardiovascular system and provided a highly useful comprehensive resource. Recent advances in single cell RNA-seq technology provide a new frontier to understand cell type specific gene expression and will allow us to further de-convolute expression patterns in cardiovascular tissues (Chaudhry et al. 2019). Further in-depth studies will be necessary to understand the gene expression networks and molecular pathways that exist in the different cardiovascular structures, and how they develop and change as the cardiovascular system matures.

## Data Availability

The datasets from the sheep gene expression atlas (Clark et al. 2017) supporting the conclusions of this article are available in the following locations. The raw read data is deposited in the European Nucleotide Archive (ENA) under study accession number PRJEB19199 (http://www.ebi.ac.uk/ena/data/view/PRJEB19199). The expression estimates (averaged across biological replicates) from Supplemental Dataset 1 can also be viewed and downloaded via BioGPS (http://biogps.org/dataset/BDS_00015/sheep-atlas/). Sample metadata for all the tissue samples collected has been deposited in the EBI BioSamples database under project identifier GSB-718 (https://www.ebi.ac.uk/biosamples/groups/SAMEG317052).

## Ethics Statement

All animal work was approved by The Roslin Institute’s Animal Welfare and Ethical Review Body (AWERB). Animals were maintained in accordance with UK Home Office guidelines and experiments were carried out under the authority of UK Home Office Project Licenses under the regulations of the Animal (Scientific Procedures) Act 1986.

## Acknowledgements

The authors would like to thank the farm staff at Dryden farm and members of the sheep tissue collection team from The Roslin Institute and R(D)SVS who were involved in tissue collections for the sheep gene expression atlas projects, particularly Ailsa Carlisle who helped to coordinate collecting the cardiovascular tissues. The authors are also grateful for the support of the FAANG Data Coordination Centre (http://data.faang.org) in the upload and archiving of the sample data and metadata and BioGPS (http://biogps.org) for hosting the sheep atlas dataset on their annotation portal.

## Authors Contributions

DAH acquired the funding for the sheep gene expression atlas. ELC coordinated and designed the sheep gene expression atlas with assistance from DAH and KMS. HGT, ELC, and KMS performed sample collection from sheep. HGT and ELC performed the RNA extractions. SJB performed all bioinformatic analyses. HGT and KMS performed the network cluster analysis. HGT performed the RT-qPCR analysis. Results were interpreted by HGT with VM, BMC and KMS. HGT wrote the manuscript with GRM, ELC, KMS and VEM. All authors read and approved the final manuscript.

## Supplementary Tables

Supplementary Table 1. Sheep primers for RT-qPCR. Primers in black were designed using Primer3 (http://primer3.ut.ee/) to span exon-exon junctions, and obtained from Invitrogen (Paisley, UK). Primers in blue were obtained from Primerdesign Ltd (Eastleigh, UK).

Supplementary Table 2. Details of tissues sequenced to generate the RNA-seq dataset for the cardiovascular gene expression atlas. Skeletal muscle (bicep) was also included, as an example of another muscle tissue, for comparative analysis. All libraries were Illumina 125 bp paired end stranded libraries.

## Supplementary Figures

Supplementary Figure 1. Gene expression profiles during development in the left ventricle. Genes include: (A) collagen type I alpha 1, *COL1A1*, (B) biglycan, *BGN*, (C) matrix metalloproteinase 2, *MMP2*, (D) TIMP metallopeptidase inhibitor 1, *TIMP1*, (E) fibrillin 1, *FBN1*, (F) fibrillin 2, *FBN2*, (G) secreted phosphoprotein1/osteopontin, *SPP1*, (H) progressive ankylosis protein, *ANKH*, (I) osteoprotegerin, *TNFRSF11B*, and (J) Runt-related transcription factor 2, *RUNX2*. Black dots show gene expression from individual animals (n = 3-5) and red dot and error bars show the mean ± standard deviation (SD) per tissue. Gene expression levels were normalised to the geomean of *GAPDH* and *YWHAZ*. In blue, asterisk (*) denotes significant differences compared to foetal d100 sheep, triangle (∆) compared to newborn sheep and circle (o) compared to 8 week old sheep, where 1 symbol = 0.01<p<0.05, 2 symbols = 0.001p<0.01 and 3 symbols = p<0.001.

Supplementary Figure 2. Gene expression profiles during development in the interventricular septum. Genes include: (A) collagen type I alpha 1, *COL1A1*, (B) biglycan, *BGN*, (C) matrix metalloproteinase 2, *MMP2*, (D) fibrillin 1, *FBN1*, (E) fibrillin 2, *FBN2*, (F) secreted phosphoprotein1/osteopontin, *SPP1*, (G) progressive ankylosis protein, *ANKH*, (H) matrix Gla protein, *MGP*, and (I) Potassium two pore domain channel subfamily K member 3, *KCNK3*. Black dots show gene expression from individual animals (n = 3-5) and red dot and error bars show the mean ± standard deviation (SD) per tissue. Gene expression levels were normalised to the geomean of *GAPDH* and *YWHAZ*. In blue, asterisk (*) denotes significant differences compared to foetal d100 sheep, triangle (∆) compared to newborn sheep, circle (o) compared to 1 week old sheep and square (□) compared to 8 week old sheep, where 1 symbol = 0.01<p<0.05, 2 symbols = 0.001<p<0.01 and 3 symbols = p<0.001.

Supplementary Figure 3. Gene expression profiles during development in the pulmonary artery. Genes include: (A) TIMP metallopeptidase inhibitor 1, *TIMP1*, (B) fibrillin 2, *FBN2*, (C) ectonucleotide pyrophosphatase/ phosphodiesterase 1, *ENPP1*, (D) secreted phosphoprotein1/osteopontin, *SPP1*, and (E) Runt-related transcription factor 2, *RUNX2*. Black dots show gene expression from individual animals (n = 3-5) and red dot and error bars show the mean ± standard deviation (SD) per tissue. Gene expression levels were normalised to the geomean of *GAPDH* and *YWHAZ*. In blue, asterisk (*) denotes significant differences compared to newborn sheep, triangle (∆) compared to 8 week old sheep, where 1 symbol = 0.01<p<0.05, 2 symbols = 0.001<p<0.01 and 3 symbols = p<0.001.

Supplementary Figure 4. Gene expression profiles during development in the aortic root. Genes include: (A) collagen type I alpha 1, *COL1A1*, (B) biglycan, *BGN*, (C) matrix metalloproteinase 2, *MMP2*, (D) fibrillin 1, *FBN1*, (E) fibrillin 2, *FBN2*, (F) ectonucleotide pyrophosphatase/ phosphodiesterase 1, *ENPP1*, (G) secreted phosphoprotein1/osteopontin, *SPP1*, and (H) Potassium two pore domain channel subfamily K member 3, *KCNK3*. Black dots show gene expression from individual animals (n = 3-5) and red dot and error bars show the mean ± standard deviation (SD) per tissue. Gene expression levels were normalised to the geomean of *GAPDH* and *YWHAZ*. In blue, asterisk (*) denotes significant differences compared to newborn sheep, triangle (∆) compared to 1 week old sheep and circle (o) compared to 8 week old sheep, where 1 symbol = 0.01<p<0.05, 2 symbols = 0.001<p<0.01 and 3 symbols = p<0.001.

Supplementary Figure 5. Gene expression profiles during development in the aortic arch. Genes include: (A) fibrillin 2, *FBN2*, (B) secreted phosphoprotein1/osteopontin, *SPP1*, (C) matrix Gla protein, *MGP*, (D) osteoprotegerin, *TNFRSF11B*, and (E) Runt-related transcription factor 2, *RUNX2*. Black dots show gene expression from individual animals (n = 3-5) and red dot and error bars show the mean ± standard deviation (SD) per tissue. Gene expression levels were normalised to the geomean of *GAPDH* and *YWHAZ*. In blue, asterisk (*) denotes significant differences compared to newborn sheep and triangle (∆) compared to 1 week old sheep, where 1 symbol = 0.01<p<0.05, 2 symbols = 0.001<p<0.01 and 3 symbols = p<0.001.

Supplementary Figure 6. Gene expression profiles during development in the abdominal aorta. Genes include: (A) collagen type I alpha 1, *COL1A1*, (B) TIMP metallopeptidase inhibitor 1, *TIMP1*, (C) fibrillin 1, *FBN1*, (D) fibrillin 2, *FBN2*, (E) secreted phosphoprotein1/osteopontin, *SPP1*, (F) progressive ankylosis protein, *ANKH*, (G) osteoprotegerin, *TNFRSF11B*, and (H) Runt-related transcription factor 2, *RUNX2*. Black dots show gene expression from individual animals (n = 3-5) and red dot and error bars show the mean ± standard deviation (SD) per tissue. Gene expression levels were normalised to the geomean of *GAPDH* and *YWHAZ*. In blue, asterisk (*) denotes significant differences compared to newborn sheep and triangle (∆) compared to 8 week old sheep, where 1 symbol = 0.01<p<0.05, 2 symbols = 0.001<p<0.01 and 3 symbols = p<0.001.

Supplementary Figure 7. mRNA expression profile for (A) matrix Gla protein (*MGP*), (B) progressive ankylosis protein homologue (*ANKH*). Gene expression levels were normalised to the geomean of *GAPDH* and *YWHAZ*. Dot plots show individual data points (black dot), the mean expression for each tissue (red dot) and standard deviation (red error bars).

Supplementary Figure 8. mRNA expression profile for (A) ecto-5’-nucleotidase (*NT5E*) and (B) Runt-related transcription factor 2 (*RUNX2*). Gene expression levels were normalised to the geomean of *GAPDH* and *YWHAZ*. Dot plots show individual data points (black dot), the mean expression for each tissue (red dot) and standard deviation (red error bars).

Supplementary Figure 9. mRNA expression profile for (A) ectonucleotide pyrophosphatase/phosphodiesterase 1 (*ENPP1*) and (B) secreted phosphoprotein1/osteopontin (*SPP1*). Gene expression levels were normalised to the geomean of *GAPDH* and *YWHAZ*. Dot plots show individual data points (black dot), the mean expression for each tissue (red dot) and standard deviation (red error bars).

Supplementary Figure 10. RNA-seq expression profiles of selected genes. Expression levels were measured using RNA-seq and shown as median expression levels in transcripts per million (TPM; n = 4-6). Y axis shows normalised median TPM (Bush et al. 2017). (A-D) Cluster 1 gene expression profiles. Genes include collagen type I alpha 1 (*COL1A1*), collagen type III alpha 1 (*COL3A1*), matrix metalloproteinase 2 (*MMP*2) and tissue inhibitor of metalloproteinases 1 (*TIMP1*). (E-G) Cluster 3 gene expression profiles. Genes include collagen type I alpha 2 (*COL1A2*), bone gamma-carboxyglutamate protein (*BGLAP*) and biglycan (*BGN*). (H-J) Cluster 22 gene expression profiles. Genes include ectonucleotide pyrophosphate/phosphodiesterase 1 (*ENPP1*), ADAM metallopeptidase with thrombospondin type 1 motif 6 (*ADAMTS6*) and SMAD family member 2 (*SMAD2*). (K-L) Cluster 24 gene expression profiles. Genes include fibrillin 1 (*FBN1*) and fibromodullin (*FMOD*). (M-O) Cluster 36 gene expression profiles. Gene include potassium two pore domain channel subfamily K member 3 (*KCNK3*), natriuretic peptide A (*NPPA*) and Dickkopf WNT signalling pathway inhibitor 3 (*DKK3*).

## Supplemental Datasets

Supplemental Dataset 1: Gene expression estimates as transcripts per million (TPM) for seven cardiovascular tissues and skeletal muscle bicep generated for the sheep gene expression atlas usin Kallisto.

Supplemental Dataset 2: Genes contained within each cluster from the gene to gene network analysis presented in Figure 1 (B) and Figure 2. Pearson correlation co-efficient r ≥ 0.99, MCL (inflation = 2.2).

