## Supplementary figures and images for "Gene expression in the cardiovascular system of the domestic sheep (*Ovis aries*); a new tool to advance our understanding of cardiovascular disease"

### Supplemental Figure 1

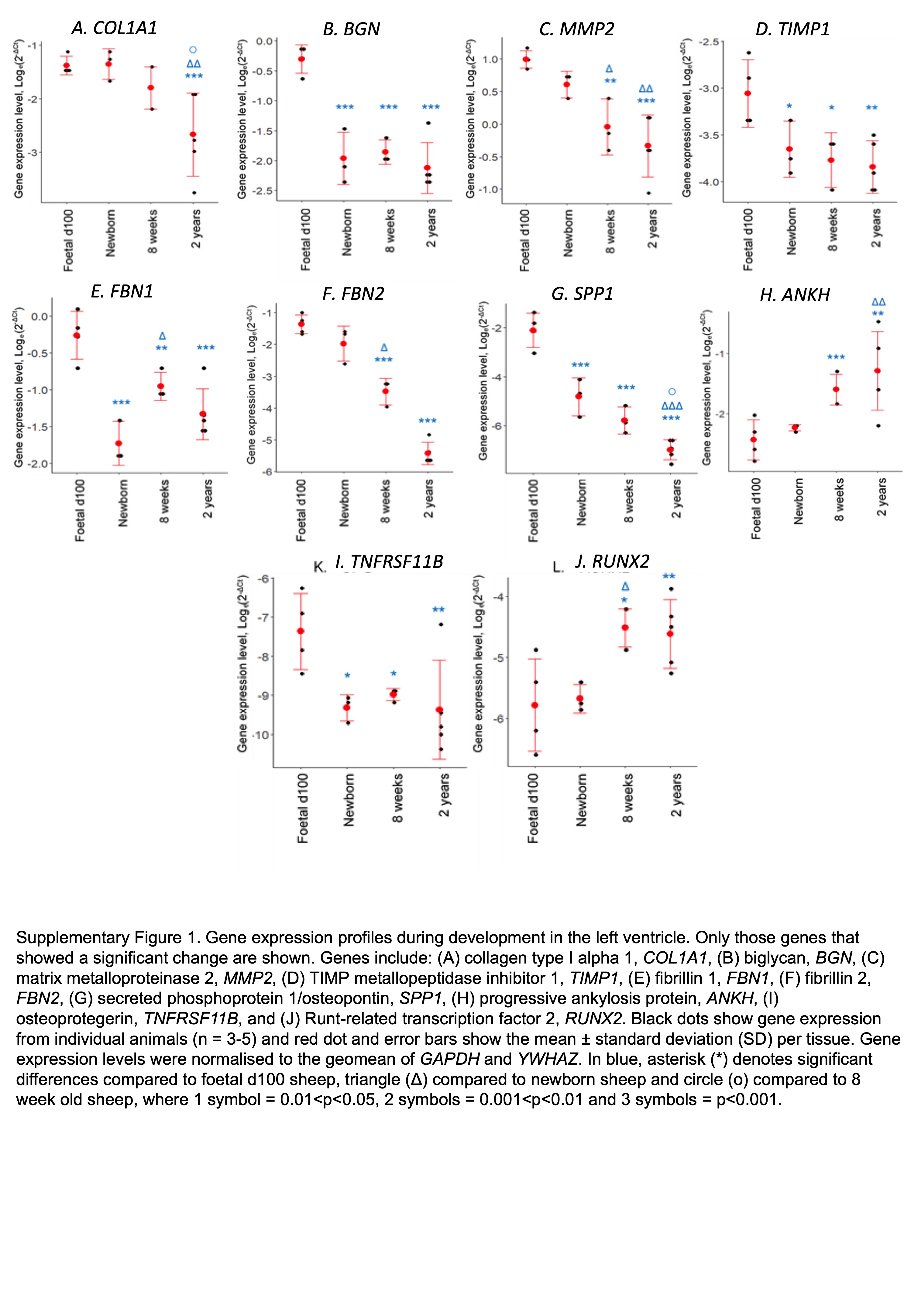

### Supplemental Figure 2

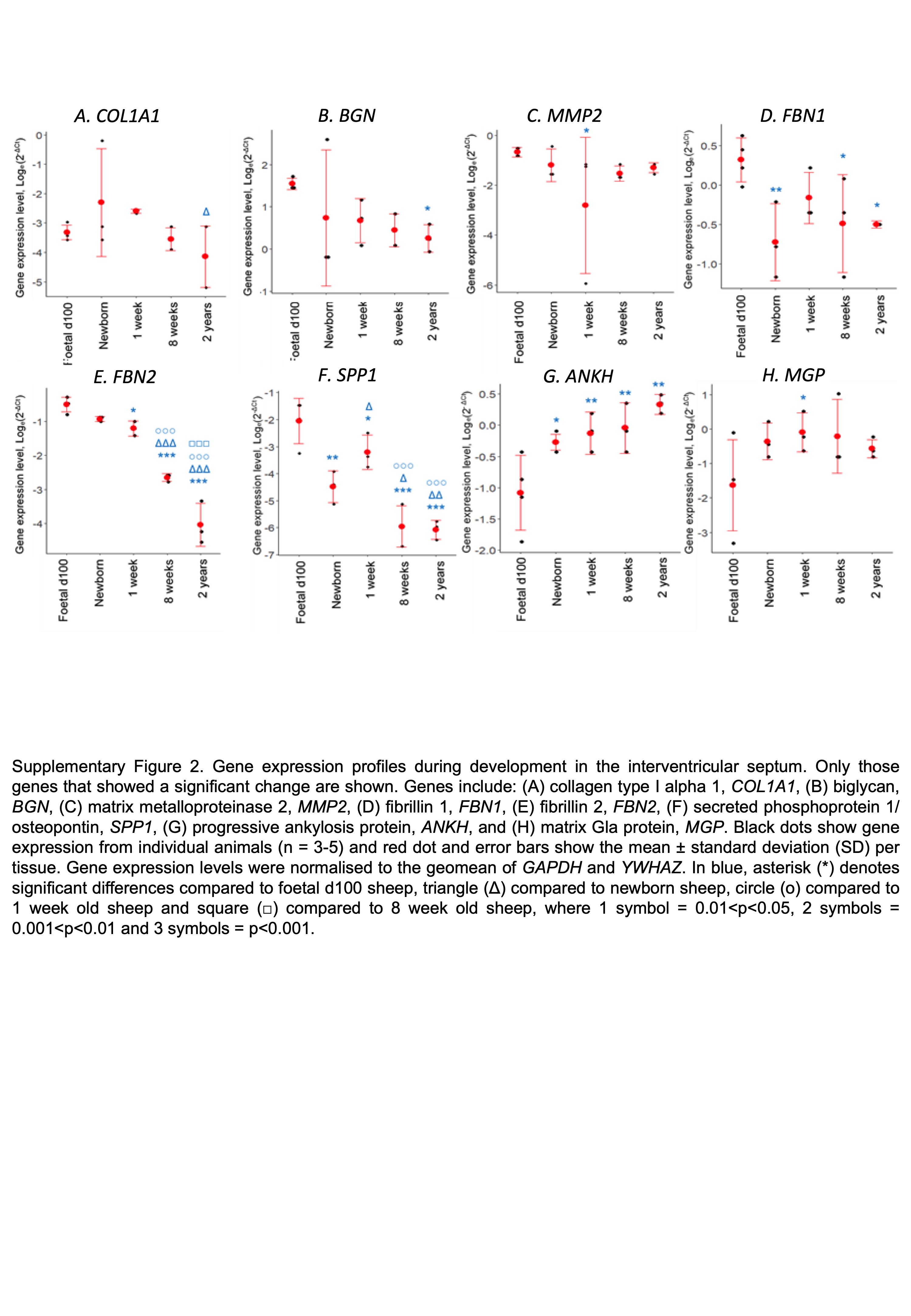

### Supplemental Figure 3

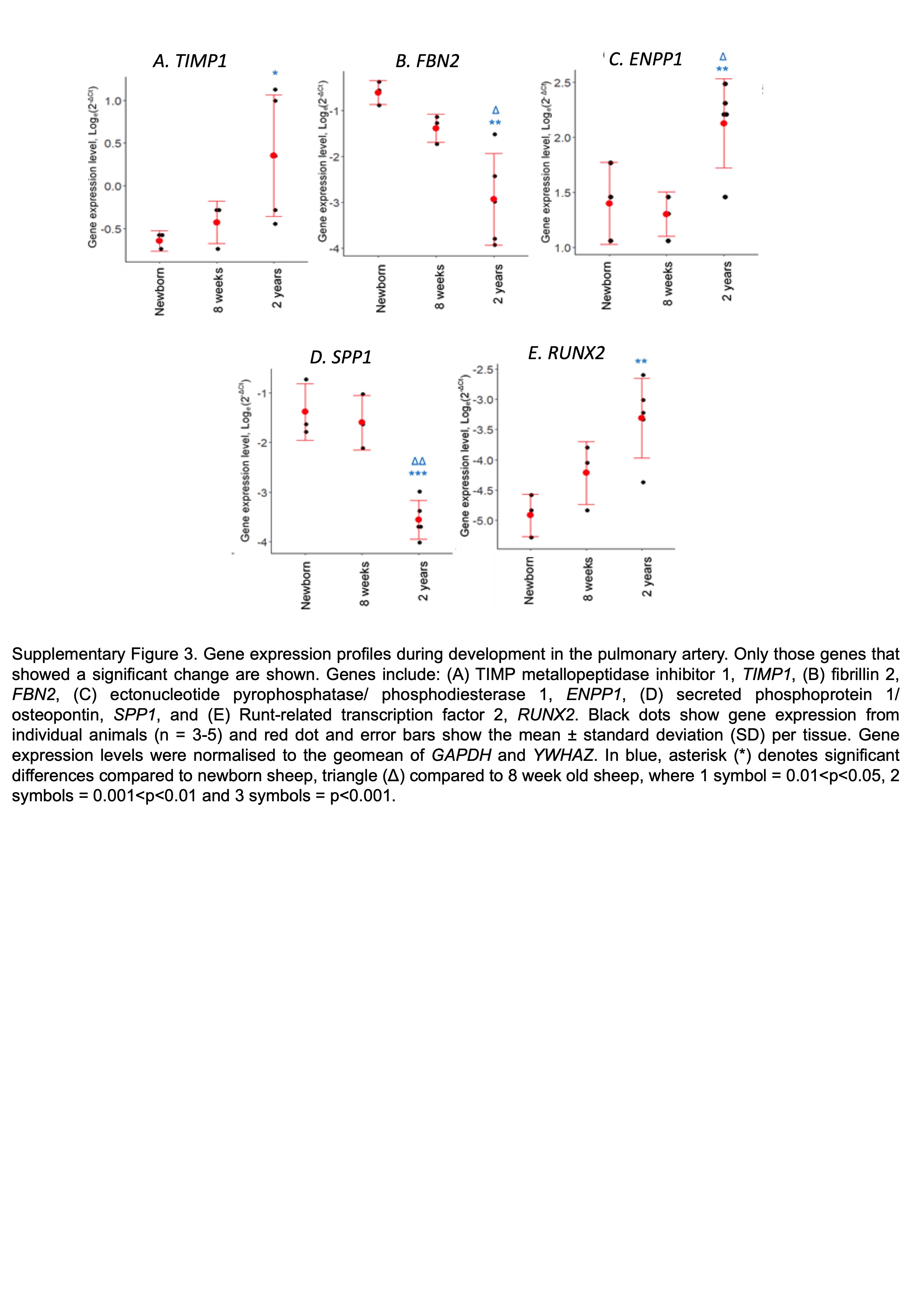

### Supplemental Figure 4

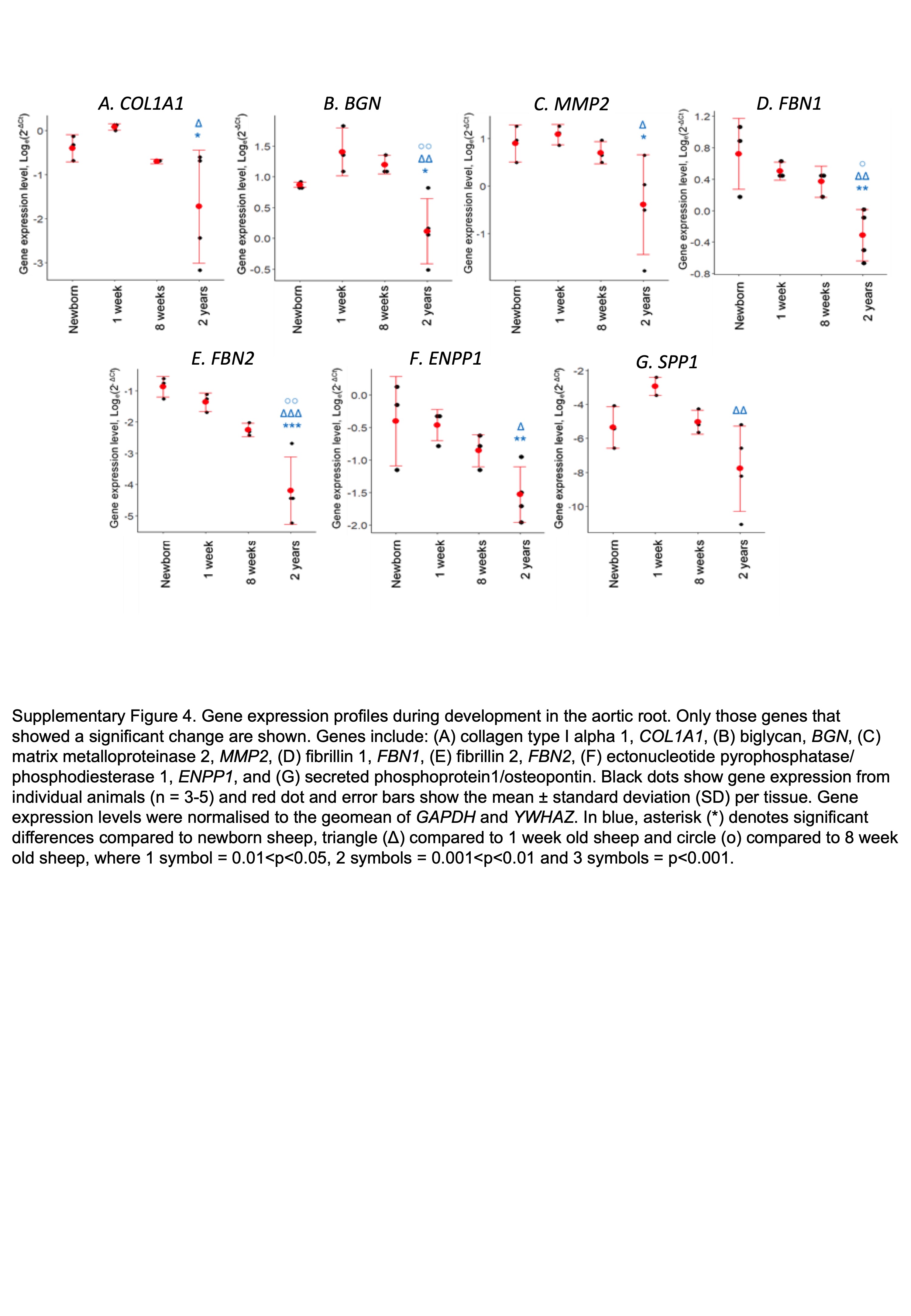

### Supplemental Figure 5

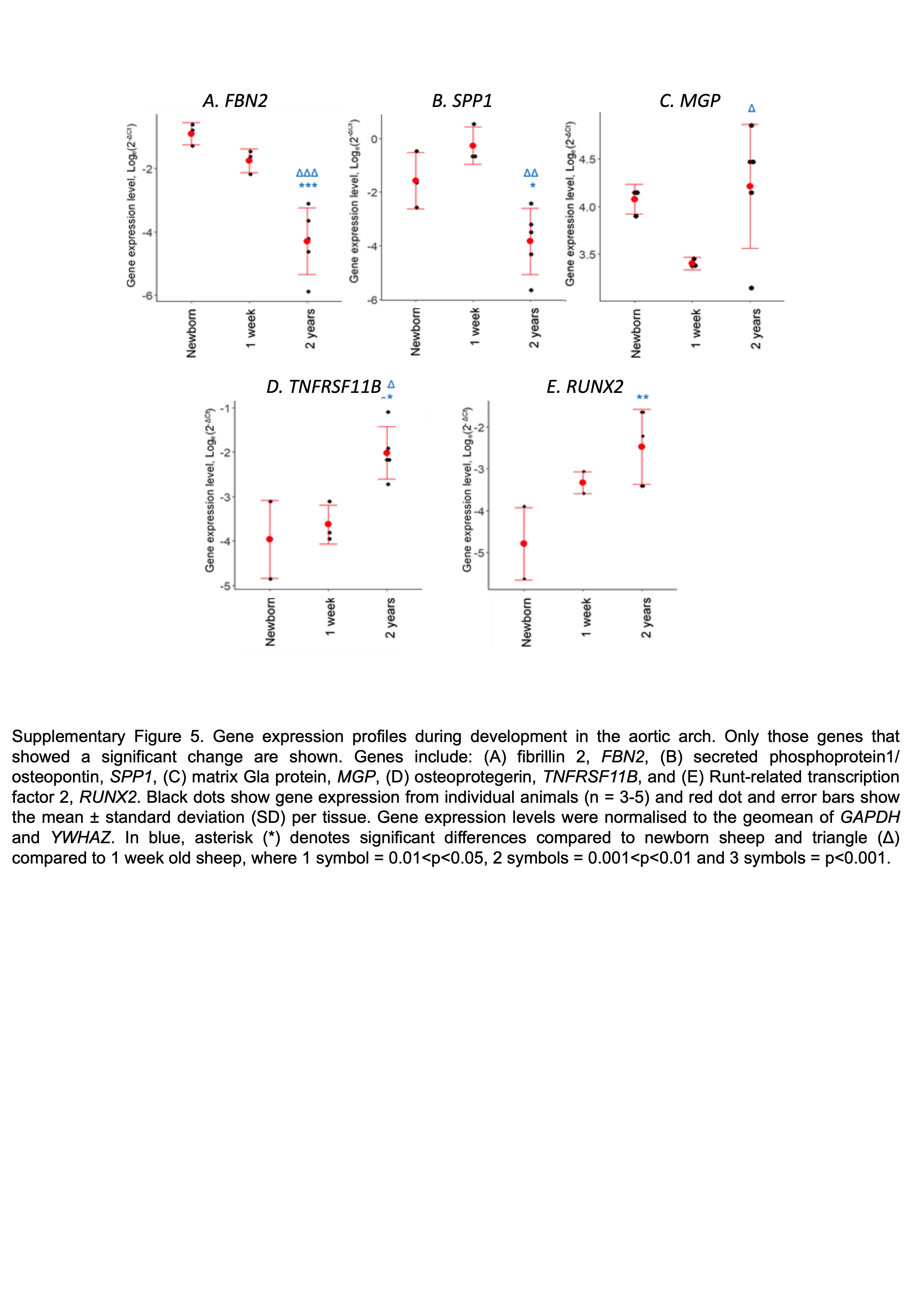

### Supplemental Figure 6

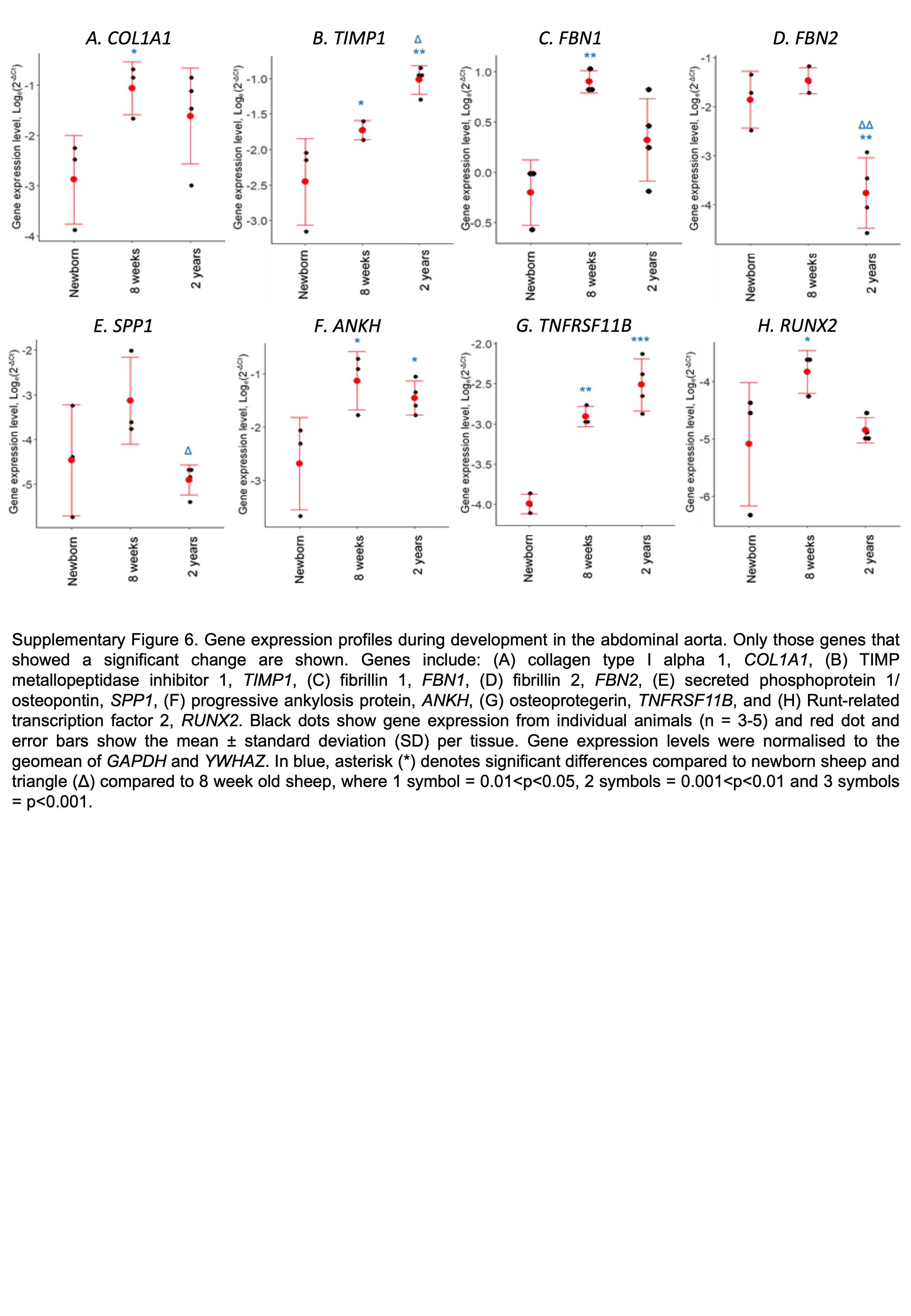

### Supplemental Figure 7

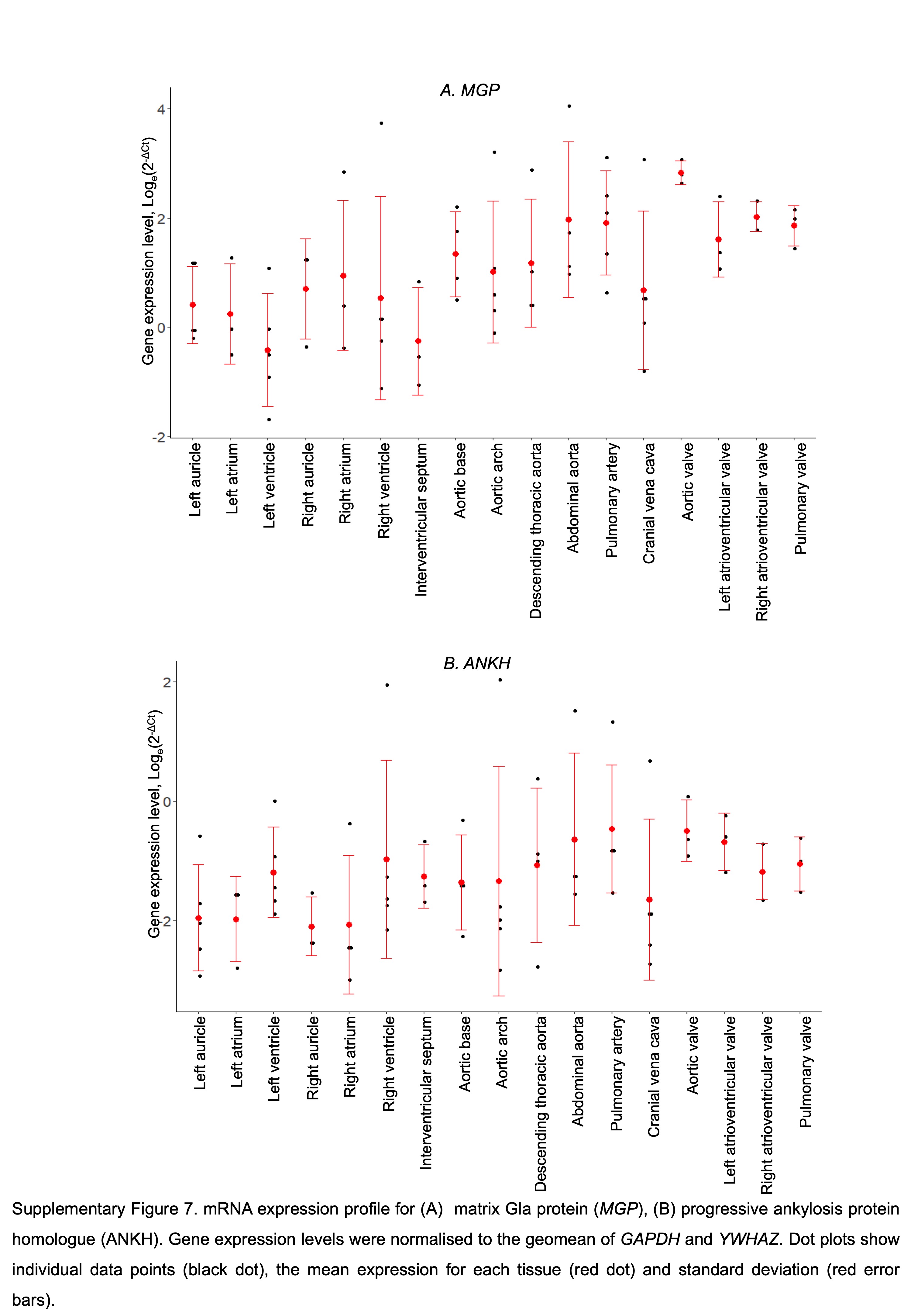

### Supplemental Figure 8

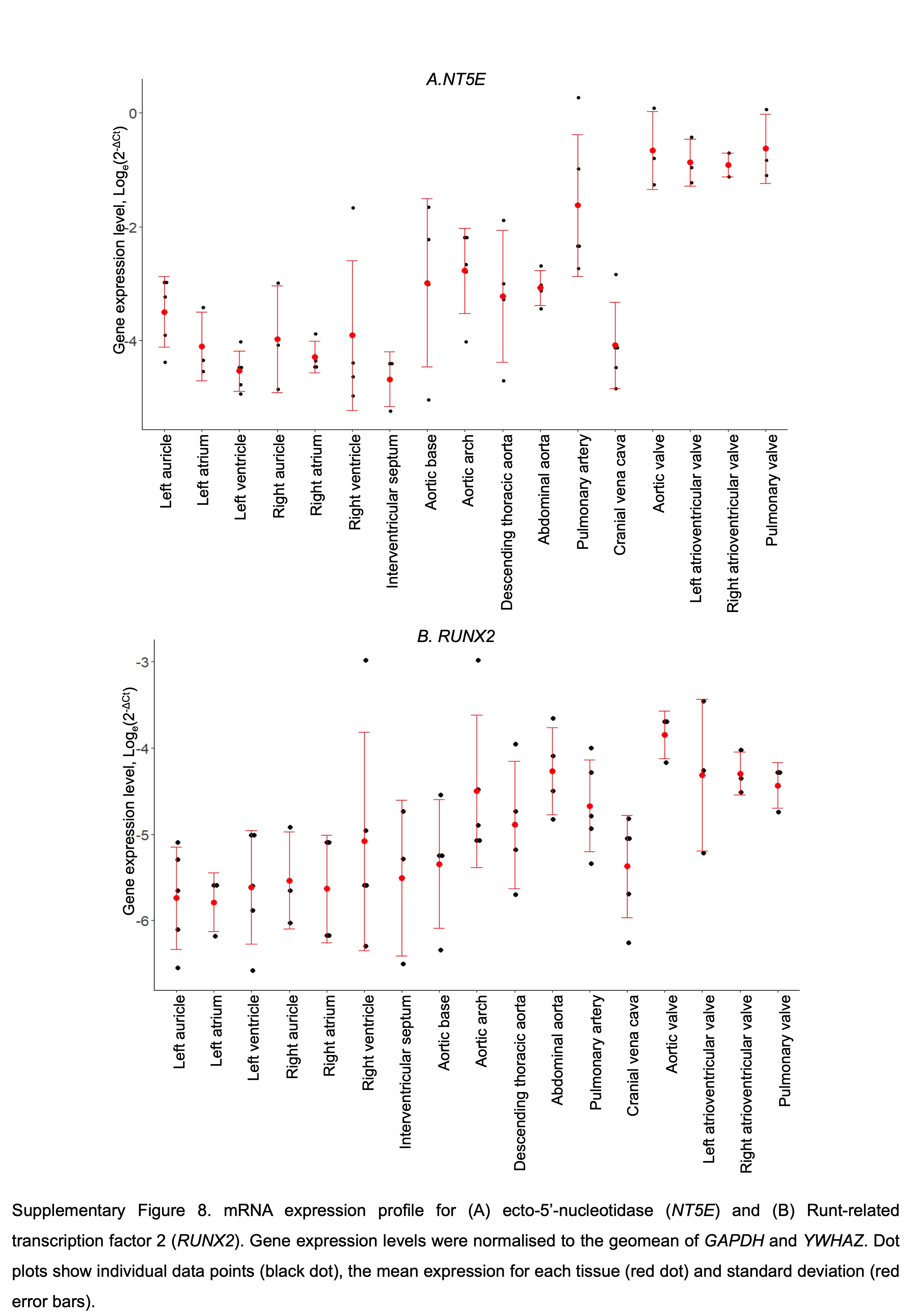

### Supplemental Figure 9

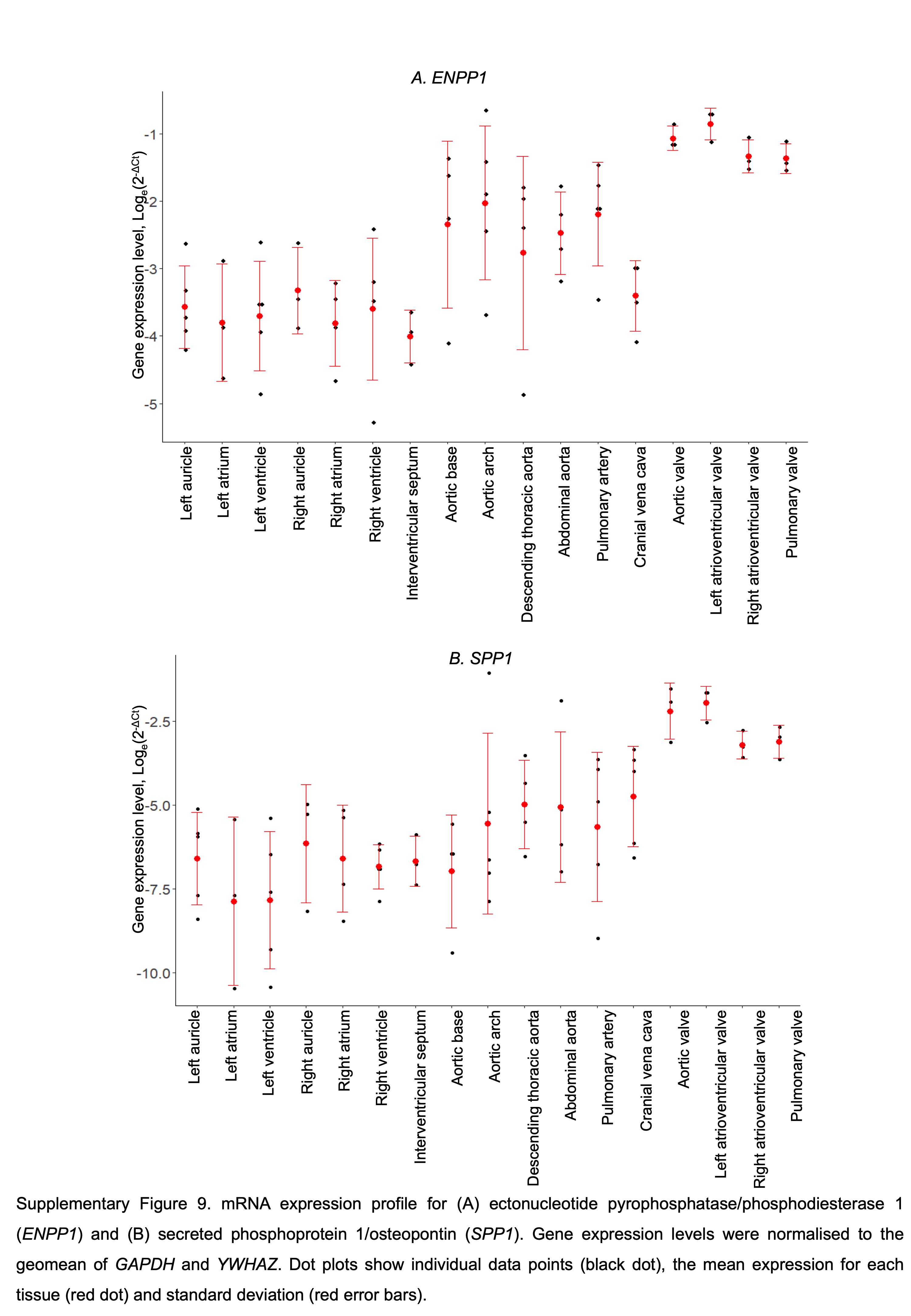

### Supplemental Figure 10

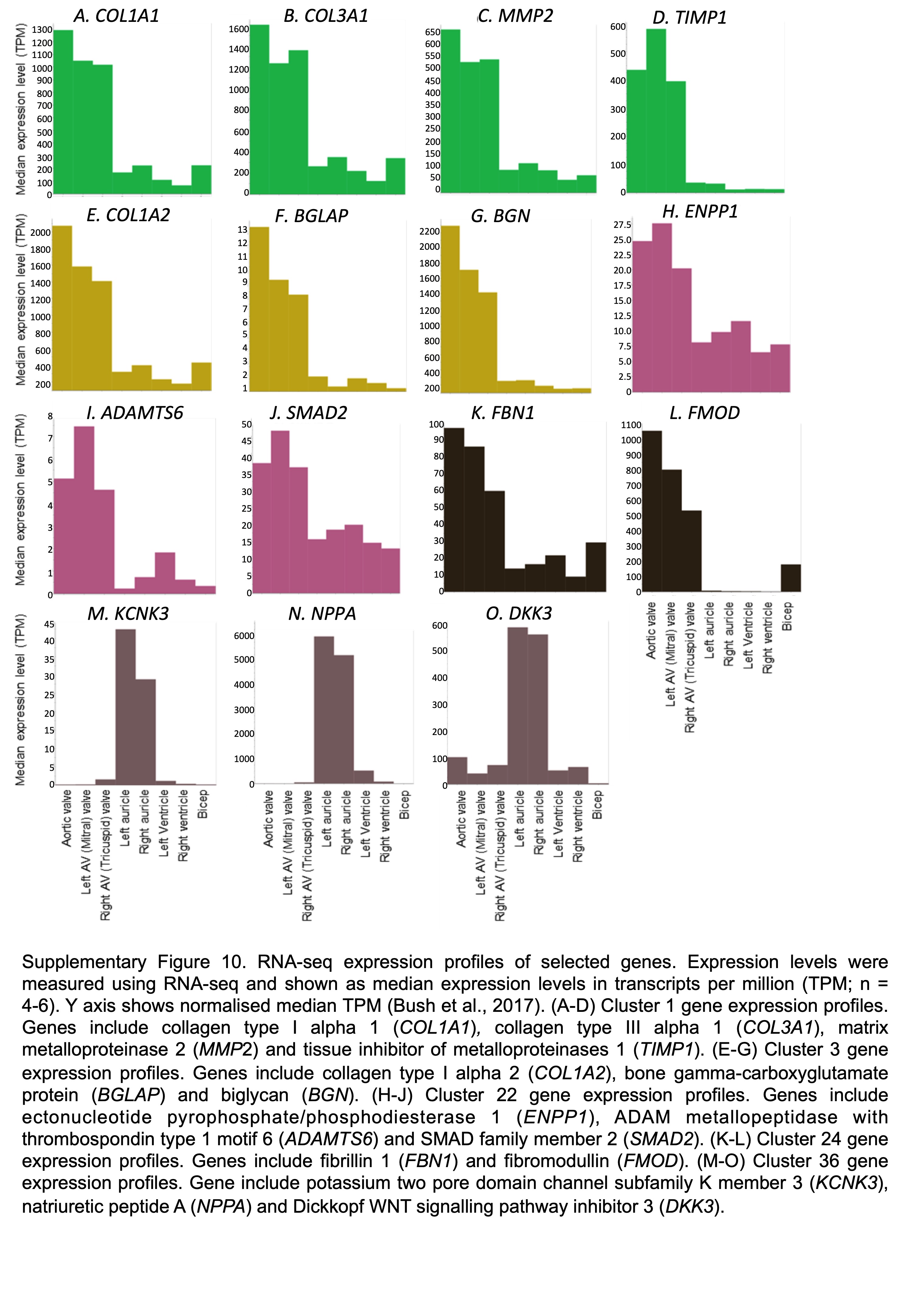

### Supplemental Table 1

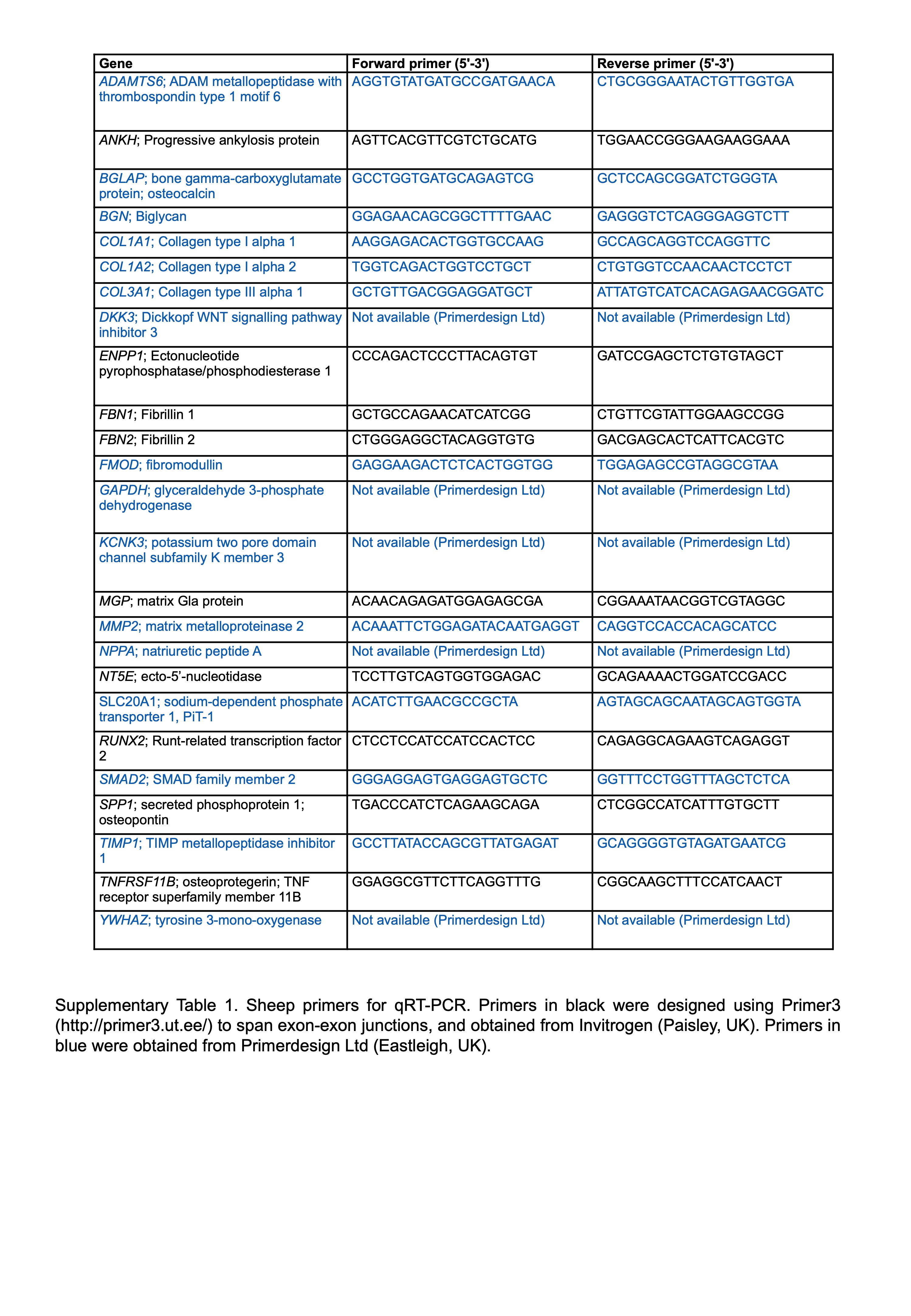

### Supplemental Table 2

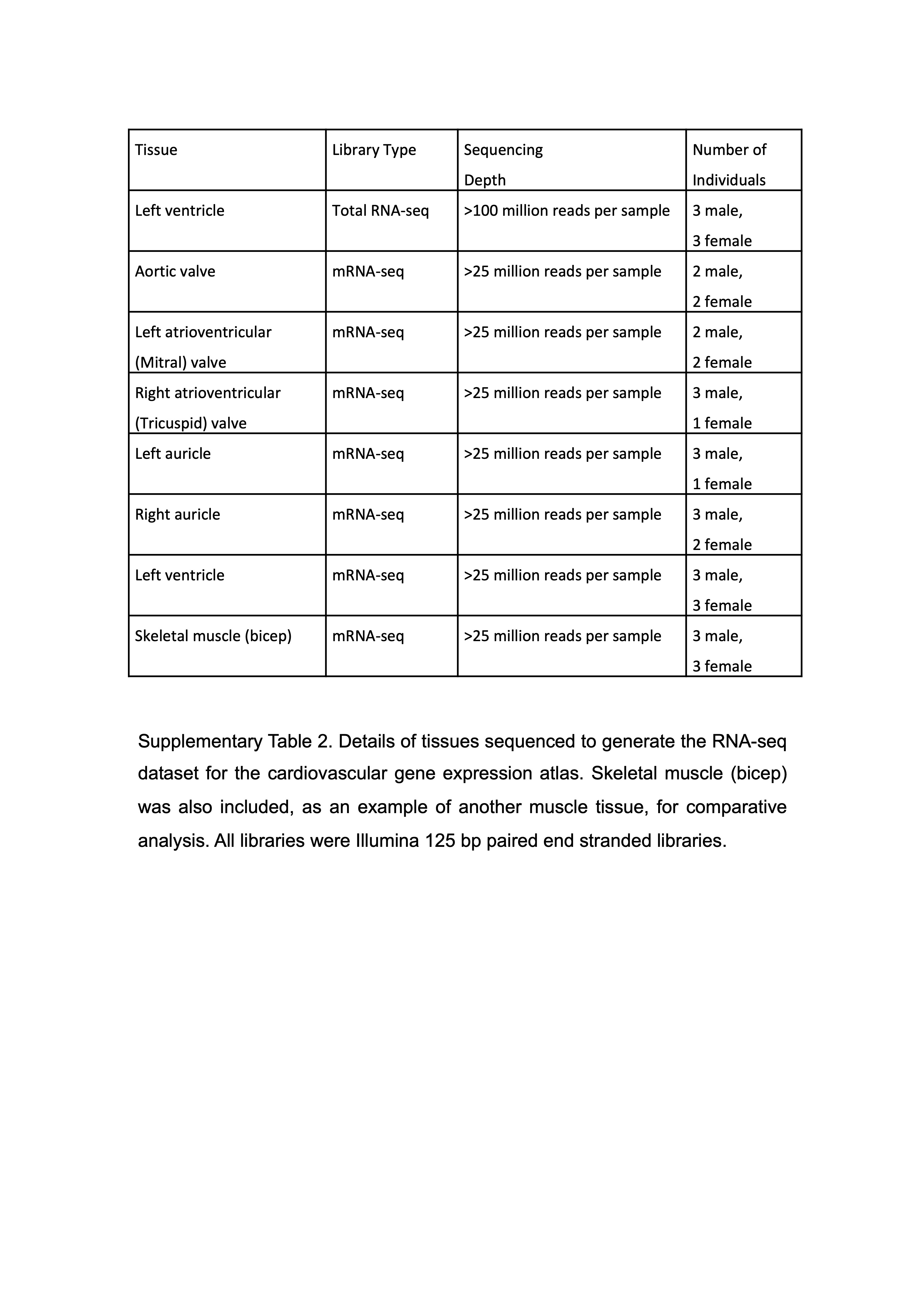

### Supplemental Table 3

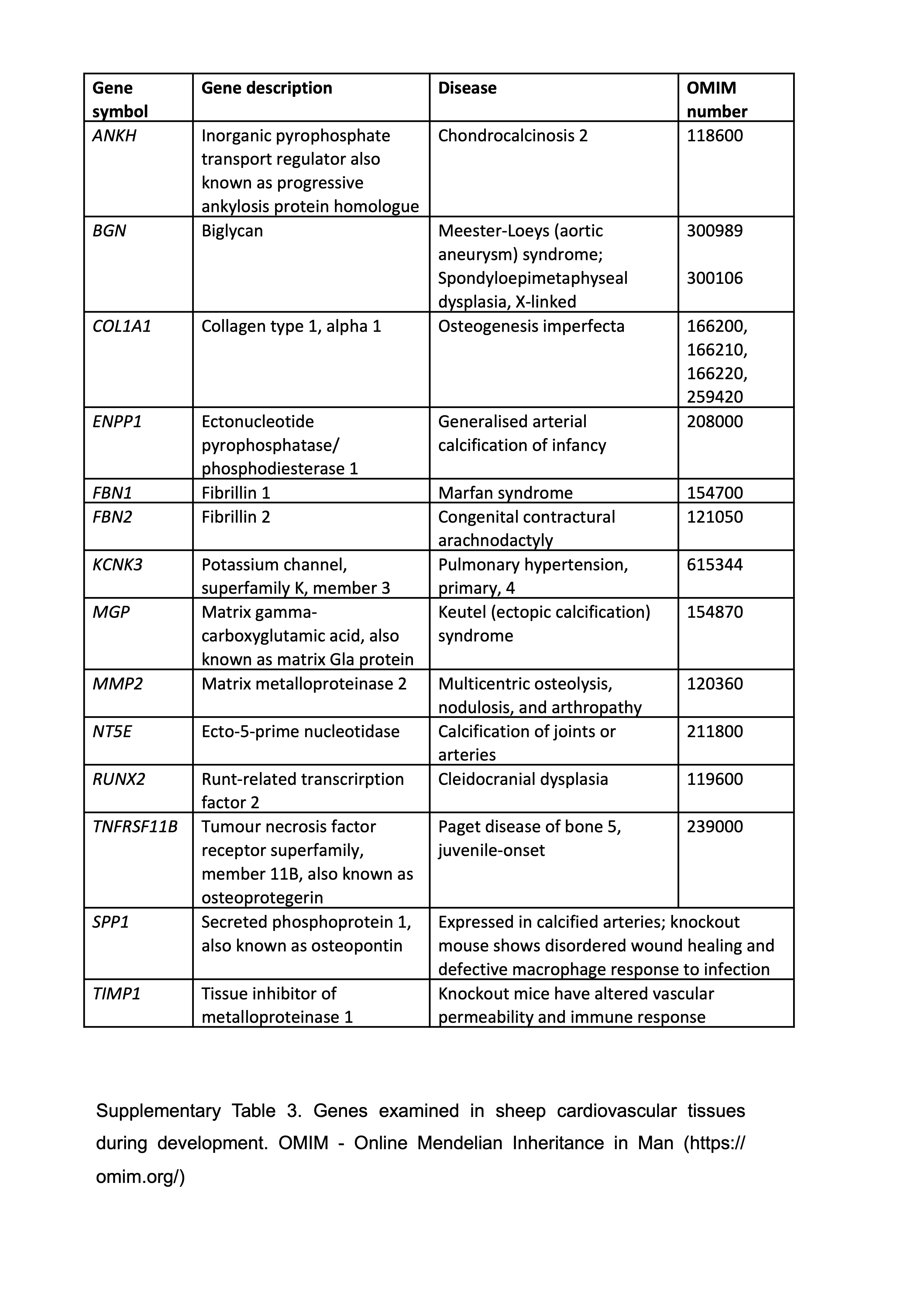
